# Association Study of Over 200,000 Subjects Detects Novel Rare Variants, Functional Elements, and Polygenic Architecture of Prostate Cancer Susceptibility

**DOI:** 10.1101/2020.02.12.929463

**Authors:** Nima C. Emami, Taylor B. Cavazos, Sara R. Rashkin, Clinton L. Cario, Rebecca E. Graff, Caroline G. Tai, Joel A. Mefford, Linda Kachuri, Eunice Wan, Simon Wong, David S. Aaronson, Joseph Presti, Laurel A. Habel, Jun Shan, Dilrini K. Ranatunga, Chun R. Chao, Nirupa R. Ghai, Eric Jorgenson, Lori C. Sakoda, Mark N. Kvale, Pui-Yan Kwok, Catherine Schaefer, Neil Risch, Thomas J. Hoffmann, Stephen K. Van Den Eeden, John S. Witte

## Abstract

The potential association between rare germline genetic variants and prostate cancer (PrCa) susceptibility has been understudied due to challenges with assessing rare variation. Furthermore, although common risk variants for PrCa have shown limited individual effect sizes, their cumulative effect may be of similar magnitude as high penetrance mutations. To identify rare variants associated with PrCa susceptibility, and better characterize the mechanisms and cumulative disease risk associated with common risk variants, we analyzed large population-based cohorts, custom genotyping microarrays, and imputation reference panels in an integrative study of PrCa genetic etiology. In particular, 11,649 men (6,196 PrCa cases, 5,453 controls) of European ancestry from the Kaiser Permanente Research Program on Genes, Environment and Health, ProHealth Study, and California Men’s Health Study were genotyped and meta-analyzed with 196,269 European-ancestry male subjects (7,917 PrCa cases, 188,352 controls) from the UK Biobank. Six novel loci were genome-wide significant in our meta-analysis, including two rare variants (minor allele frequency < 0.01, at 3p21.31 and 8p12). Gene-based rare variant tests implicated a previously discovered PrCa gene (*HOXB13*) as well as a novel candidate (*ILDR1*) highly expressed in prostate tissue. Haplotypic patterns of long-range linkage disequilibrium were observed for rare genetic variants at *HOXB13* and other loci, reflecting their evolutionary history. Furthermore, a polygenic risk score (PRS) of 187 known, largely common PrCa variants was strongly associated with risk in non-Hispanic whites (90^th^ vs. 10^th^ decile OR = 7.66, *P* = 1.80*10^-239^). Many of the 187 variants exhibited functional signatures of gene expression regulation or transcription factor binding, including a six-fold difference in log-probability of Androgen Receptor binding at the variant rs2680708 (17q22). Our finding of two novel rare variants associated with PrCa should motivate further consideration of the role of low frequency polymorphisms in PrCa, while the considerable effect of PrCa PRS profiles should prompt discussion of their role in clinical practice.

## INTRODUCTION

For a number of diseases, including prostate cancer (PrCa), there has been limited success in detecting associated rare genetic variants, some of which may have substantial effect sizes [1]. This is in part due to the difficulty of measuring or imputing rare variants in adequately powered studies. Still, some rare germline variants associated with prostate cancer have been detected, such as in the DNA damage repair gene *BRCA2* [2] and the developmental transcription factor *HOXB13* [3]. While relatively few rare variants have been discovered, in aggregate they may comprise a substantial portion of PrCa risk heritability [4]. In contrast, genome-wide association studies (GWAS) of more common variants have identified over 150 independent genetic variants associated with PrCa [5]. Each variant is typically associated with only a modest increase in PrCa risk, and thus not of sufficient magnitude to be clinically significant. However, combining all associated variants together into a single polygenic risk score (PRS) may distinguish men with a meaningfully increased risk of PrCa.

To investigate the impact of rare and common variants on PrCa, we undertook a large scale genome-wide study of over 200,000 male subjects from two large cohorts: Kaiser Permanente (KP) in California [6] and the UK Biobank (UKB) [7]. Genotype microarrays, including GWAS backbones and custom rare variant content, were assayed in both cohorts, and unmeasured genotypes were imputed using a reference panel of over 27,000 phased Haplotype Reference Consortium (HRC) genomes [8]. We evaluated associations between individual rare and common variants and PrCa risk and interpreted the evolutionary origin and functional mechanisms of novel findings using multi-omics data. We also performed PRS modeling and functional characterization for the known common risk variants.

## METHODS AND MATERIALS

### Study Populations

We studied two cohorts of PrCa cases and non-diseased controls: 1) KP subjects from the Northern California Research Program on Genes, Environment and Health (RPGEH), the California Men’s Health Study (CMHS) and the ProHealth Study; and 2) the UKB. The KP cohort included 6,196 male cases and 5,453 male controls of European-ancestry (mean age at diagnosis for cases = 68.1 years, mean age at baseline among controls = 71.5). The UKB cohort included 7,917 cases and 188,352 controls of European ancestry (mean age at diagnosis = 64.1, mean age among controls = 57.1). Subject demographics and characteristics are described in detail in Supplementary Table 1.

### Custom Microarray Design and Genotyping

To directly assay or tag putatively functional rare variation in samples from KP, we collaborated with Affymetrix Inc. on the design of a custom Axiom DNA microarray (Supplementary Figure 1a) that was complementary to the GWAS array previously genotyped in the KP population [9]. The algorithm used to select variants on the custom array (Supplementary Figure 1b) resulted in 416,047 variant probesets comprising 54 distinct modules, including missense and loss-of-function mutations, rare exonic mutations from The Cancer Genome Atlas (TCGA) and dbGaP prostate cancer tumor exomes [10, 11], and variants to supplement the previously genotyped GWAS array [6] (Table 2). Many modules and most of the design content overlapped with the probesets on the UKB Affymetrix Axiom array, for which the array design, sample processing, and genotyping have been detailed [7].

Saliva biospecimens from KP participants were processed for DNA extraction using a protocol previously reported [9]. DNA samples from KP were processed using Samasy [12], a sample management system providing a visual and machine interface to facilitate robot liquid handling automation from source plates to destination plates matched by age, case status, and ethnicity. The algorithm implemented for destination plate randomization is described in the Supplementary Materials. A total of 173 96-well destination plates were amplified to increase DNA yields, and 200 ng of input DNA per well were array hybridized for 48 hours at 48 °C and genotyped using an Affymetrix GeneTitan Multi-Channel instrument.

### Quality Control and Imputation

Detailed descriptions of the sample and genotype quality control (QC) procedures are given in the Supplementary Materials. Briefly, for the KP samples, we excluded specimens with poor resolution fluorescent measurements (DQC < 0.75) or call rate < 0.95 (Supplementary Figure 2a). Based on heterozygosity rate, call rate, and plate call rate, samples were further stratified into three tiers that were used to guide genotype quality control. Specifically, genotype calls and posterior cluster locations from higher tier samples (as a consequence of higher input DNA quantities) were prioritized and used as empirical priors for resolving genotypes of lower tier samples using the Affymetrix AxiomGT1 algorithm (Supplementary Figure 2b) [13]. Genotypes were also filtered based on batch differences across the RPGEH, CMHS, and ProHealth, and based on the fold-difference in minor allele frequency (MAF) relative to the HRC and 1000 Genomes Project reference panels. These genotypes were then merged with previously assayed GWAS genotypes for the KP subjects, whose QC was described in a prior publication [6].

The KP data were phased using Eagle v2.3 (cohort-based) [14], and imputed using Minimac3 to two reference panels: (1) a subpopulation of 27,165 HRC genomes accessible via the European Genome Archive (EGAS00001001710, which includes the 1000 Genomes Project Phase III samples), and (2) the 1000 Genomes Project Phase III reference panel (2,514 genomes). Single nucleotide variant calls were imputed using the union of (1) and (2), and indel polymorphisms were imputed using (2) (not yet part of the HRC due to additional difficulty in harmonizing indels; Supplementary Figure 3). Variants with r^2^_INFO_ < 0.3 and with a minor allele frequency less than 1/N_REF_, where N_REF_ represents the total number of chromosomes in the reference panel, were removed from the imputed genotypes. Individuals were ultimately classified into ethnic analysis groups (African, East Asian, European, or Hispanic ancestry) based on self-reported ethnicity [15, 16], although only European ancestry subjects were retained for this study due to the sample size necessary to detect rare genetic variant associations.

For the UKB data, pre-imputation QC protocols have been previously described [7]. Genotypes were imputed using two reference panels: the complete HRC reference (64,976 haplotypes) [8], and the combined UK10K plus 1000 Genomes Project Phase III reference panels (9,746 haplotypes). We similarly excluded poorly imputed (*r*^2^_INFO_ < 0.3) and excessively rare (MAF < 3*10^-5^) genotypes from the UKB.

### Association Analyses

Associations between variant genotypes and prostate cancer were evaluated for European-ancestry subjects using logistic regression with adjustment for age (for PrCa cases, age at diagnosis, versus age at time of study enrollment for controls), body mass index, genotyping array, and principal components of ancestry (PCs). The KP models controlled for 20 PCs using PLINK v2.00 [17], and the UKB models were adjusted for 10 PCs. The KP and UKB data were combined by fixed-effect meta-analysis using Metasoft v2.0.0 [18]. Gene-based rare variant tests (observed MAF < 1%) were conducted with the Sequence Kernel Association Test (SKAT) using the rvtests package (v20171009) [19], and meta-analyzed by Fisher’s method [20] using R v3.3.3.

### Evolutionary History of Rare Variants

To quantify the recency in origin of rare prostate cancer risk variants, we examined the extended haplotype homozygosity (EHH), or the length of a haplotype on which a variant allele resides, using the reference panel of 27,165 phased HRC genomes and the selscan package [21]. We also quantified the integrative haplotype score (iHS), or log ratio between a variant’s major and minor alleles of the area under the EHH curves for each allele [21], to reflect differences in allelic age or selective pressure between the derived and ancestral alleles. The iHS was computed using an EHH cutoff of 0.05, including both upstream (iHS_L_) and downstream (iHS_R_) of the query position.

### Polygenic Risk Score Analyses

For each individual, their PRS was computed by multiplying the out-of-sample effect sizes [5, 6] for each of the 187 previously reported PrCa risk loci (log ORs) by their genotype dosages, and then summing the resulting 187 values together (Supplementary Table 3). The odds ratios and 95% confidence intervals for associations between standardized PRS values (mean = 0, standard deviation = 1) and prostate cancer case-control status were estimated using logistic regression with adjustment for the same covariates modeled in our association analyses, with the exception of genotyping array so they could be compared.

### Functional Annotation

To consider the functional relevance of the known PrCa risk variants, we integrated two different analyses and sources of data. We trained elastic net regression models of normal prostatic gene expression [22], with a linear combination of germline genotypes as the predictor, using GLMNet [23] and a dataset of 471 subjects with normal prostate tissue RNA expression and genotype data [24]. Among the 187 previously reported prostate cancer risk variants, as well as the novel genome-wide significant variants identified here, those directly modeled or in linkage disequilibrium (LD *r*^2^ > 0.5) with a modeled variant in our expression models were reported. For the same set of variants, allele-specific differential transcription factor binding affinity was also estimated using sTRAP transcription factor affinity prediction [25] with the major and minor alleles.

## RESULTS

### Variant Association Analysis and Evolutionary Characterization

Genome-wide significant associations (*P*_Meta_ < 5*10^-8^) were observed at six novel loci (>3 Mb away and LD *r*^2^ < 0.005 in all 1000 Genomes Phase III populations, relative to known loci). Among the six loci (Figure 1; Table 1), three variants (rs557046152, rs555778703, and rs62262671) were at least nominally significant with consistent directions of effect in both the KP and UKB data, and two of these were rare imputed variants in European ancestry populations: rs557046152 (MAF = 0.003) and rs555778703 (MAF = 0.009). The remaining three variants were associated only in the UK Biobank. An additional gene-based rare variant meta-analysis of KP and UKB, using the sequence kernel association test (SKAT) and variants with MAF < 0.01, yielded a significant association at *HOXB13* (*P* = 1.72*10^-7^; Supplementary Figure 4), a well-characterized prostate cancer risk locus harboring a rare yet highly penetrant missense founder mutation rs138213197 [3]. SKAT also identified a suggestive *P*-value for *ILDR1* (*P* = 7.46*10^-6^), a gene primarily expressed in prostate tissue [26].

**Figure 1.**
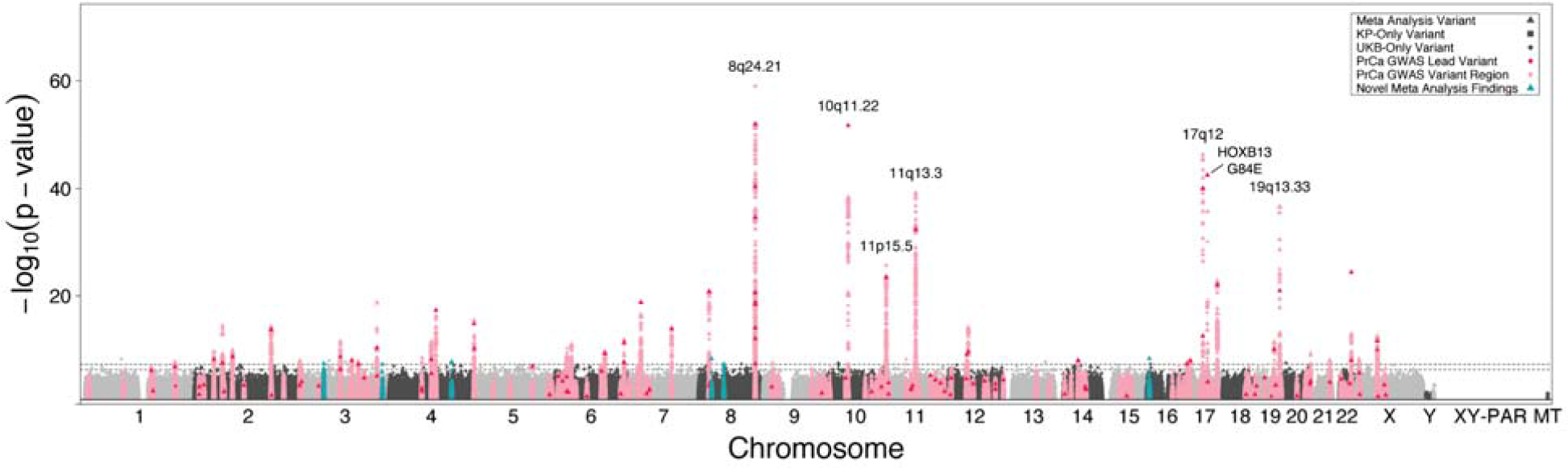
Prostate Cancer Risk Meta-Analysis Manhattan Plot for Kaiser Permanente and UK Biobank European-Ancestry Subjects. Genome-Wide Manhattan Plot of Prostate Cancer Risk. Manhattan plot depicting the results of a meta-analysis of male European-ancestry subjects from the Kaiser Permanente (KP; N = 6,196 PrCa cases, 5,453 controls) and UK Biobank (UKB; N = 7,917 PrCa cases, 188,352 controls) cohort genome-wide associations with prostate cancer (PrCa) risk. The associations (-log_10_(*P*-value), Y-axis) are plotted against the chromosome (1-22, X, Y, XY-pseudoautosomal region XY-PAR, and mitochondrial chromosome MT) and position (X-axis) of the genotyped or imputed genetic variants, with thresholds for significant (*P* < 5.0*10^-8^) and suggestive (5.0*10^-7^ < *P* < 5.0*10^-8^) associations illustrated by dashed grey lines. Non-significant loci on odd and even chromosomes are colored in alternating shades, and all variants with *P* > 0.05 are excluded from the plot. Triangular data points illustrate variants that were meta-analyzed between KP and UKB, while squares and circles indicate variants present exclusively in the KP or UKB summary statistics, respectively. Previously discovered PrCa loci are highlighted in pink for a 2 Mb window around the reported lead variant, which is highlighted in red, and previously unreported loci reaching genome-wide significance in our meta-analysis are colored in teal.

**Table 1.**
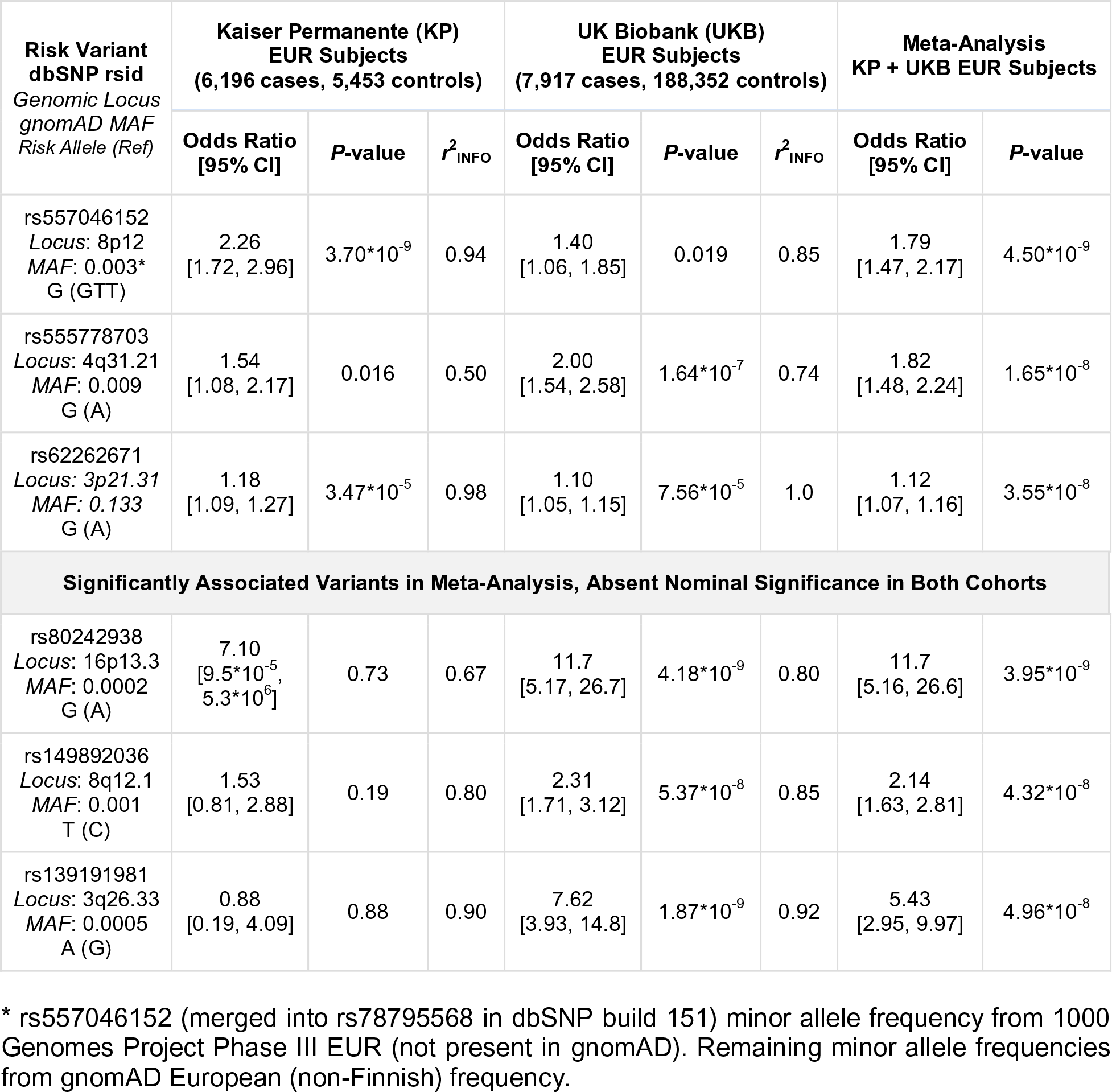
Novel Prostate Cancer Susceptibility Associations from the Meta-Analysis of European Ancestry Subjects from Kaiser Permanente and UK Biobank

**Table 2.** Normal Prostate Tissue Expression eQTLs Correlated with PrCa Risk Variants

| Previously Reported PrCa Risk Variant | Gene Name | Number of eQTL Variants Targeting Gene and with LD $r^2 > 0.5$ with Risk Variant | eQTL Variants (chr.hg19pos.ref.alt) |
| --- | --- | --- | --- |
| rs17599629 | LYSMD1 | 1 eQTLs | rs17599629 (chr1.150658287.A.G) |
| rs1775148 | RAB7L1 | 1 eQTLs | rs1775148 (chr1.205757824.C.T) |
| rs13385191 | C2orf43 | 1 eQTLs | rs13385191 (chr2.20888265.A.G) |
| rs2430386 | EHBP1 | 3 eQTLs | rs201697978 (chr2.62876580.A.C),<br>rs12713462 (chr2.62804482.C.T),<br>rs142973842 (chr2.63056706.TTG.T) |
| rs13016083 | ACVR2A | 4 eQTLs | rs7423878 (chr2.148689369.T.C),<br>rs7600869 (chr2.148551232.C.G),<br>rs70992173 (chr2.148570502.AT.A),<br>rs1424949 (chr2.148542963.T.G) |
| rs62262671 | RBM6 | 1 eQTLs | rs62262671 (chr3.49649873.A.G) |
|  | UBA7 | 1 eQTLs | rs62262671 (chr3.49649873.A.G) |
| rs12653946 | IRX4 | 1 eQTLs | rs12653946 (chr5.1895829.C.T) |
| rs1983891 | FOXP4 | 5 eQTLs | rs913074 (chr6.41538545.T.C),<br>rs4714486 (chr6.41542417.C.T),<br>rs4714485 (chr6.41536587.T.G),<br>rs1886816 (chr6.41544494.A.G),<br>rs6458228 (chr6.41543793.C.A) |
| rs9469899 | UHRF1BP1 | 1 eQTLs | rs9469899 (chr6.34793124.G.A) |
| rs1933488 | RGS17 | 1 eQTLs | rs6557267 (chr6.153433701.C.T) |
| rs9364554 | SLC22A3 | 2 eQTLs | rs1112444 (chr6.160835192.C.A),<br>rs9364554 (chr6.160833664.C.T) |
| rs6465657 | BHLHA15 | 1 eQTLs | rs6465657 (chr7.97816327.C.T) |
| rs1182 | C9orf78 | 5 eQTLs | rs55946414 (chr9.132583289.A.T),<br>rs1043186 (chr9.132573290.C.T),<br>rs3842225 (chr9.132575426.GC.G),<br>rs11787741 (chr9.132578284.A.G),<br>rs13283469 (chr9.132582014.C.T) |
| rs10993994 | MSMB | 1 eQTLs | rs10993994 (chr10.51549496.T.C) |
|  | NCOA4 | 1 eQTLs | rs10993994 (chr10.51549496.T.C) |
|  | AGAP7 | 1 eQTLs | rs10993994 (chr10.51549496.T.C) |
| rs4962416 | CTBP2 | 5 eQTLs | rs4962416 (chr10.126696872.T.C),<br>rs12769019 (chr10.126697327.A.G),<br>rs4962720 (chr10.126696840.G.T),<br>rs12769682 (chr10.126697494.G.C),<br>rs4962419 (chr10.126697114.G.A) |
| rs61890184 | PPFIBP2 | 1 eQTLs | rs61890184 (chr11.7547587.G.A) |
| rs12785905 | SYT12 | 2 eQTLs | rs12785905 (chr11.66951965.G.C),<br>rs12785906 (chr11.66951966.G.C) |
| rs11568818 | MMP7 | 1 eQTLs | rs11568818 (chr11.102401661.T.C) |
| rs11214775 | TMPRSS5 | 1 eQTLs | rs11214775 (chr11.113807181.G.A) |
| rs138466039 | PKNOX2 | 1 eQTLs | rs138466039 (chr11.125054793.C.T) |
| rs80130819 | COL2A1 | 1 eQTLs | rs80130819 (chr12.48419618.A.C) |
| rs684232 | FAM57A | 2 eQTLs | rs2474694 (chr17.618039.G.A),<br>rs684232 (chr17.618965.T.C) |
|  | GEMIN4 | 2 eQTLs | rs2474694 (chr17.618039.G.A),<br>rs684232 (chr17.618965.T.C) |
| rs142444269 | C17orf79 | 1 eQTLs | rs142444269 (chr17.30098749.C.T) |
| rs12956892 | SEC11C | 16 eQTLs | rs4940816 (chr18.56745159.A.G),<br>rs4940817 (chr18.56745263.T.G),<br>rs4940815 (chr18.56745144.A.G),<br>rs4940812 (chr18.56742965.G.A),<br>rs4940810 (chr18.56742446.T.C),<br>rs4940811 (chr18.56742904.A.G),<br>rs12956892 (chr18.56746315.G.T),<br>rs12327532 (chr18.56744666.T.G),<br>rs12326997 (chr18.56743138.A.G),<br>rs12327517 (chr18.56744475.T.C),<br>rs12327515 (chr18.56744457.T.C),<br>s10579935 (chr18.56742710.GTAAA.G),<br>rs12327308 (chr18.56743208.T.G),<br>rs4940809 (chr18.56742291.T.C),<br>rs34192989 (chr18.56744092.T.C),<br>rs4940442 (chr18.56742873.A.G) |
| rs7241993 | ATP9B | 1 eQTLs | rs9967549 (chr18.76774276.A.C) |
| rs8102476 | CATSPERG | 2 eQTLs | rs8102476 (chr19.38735613.C.T),<br>rs8102454 (chr19.38735480.G.A) |
|  | PPP1R14A | 2 eQTLs | rs8102476 (chr19.38735613.C.T),<br>rs8102454 (chr19.38735480.G.A) |
| rs5945572 | NUDT11 | 5 eQTLs | rs1327304 (chrX.51214176.C.A),<br>rs1327302 (chrX.51210615.G.A),<br>rs5945572 (chrX.51229683.A.G),<br>rs58498379 (chrX.51223415.C.T),<br>rs1327303 (chrX.51214169.C.T) |
| rs4844289 | NLGN3 | 1 eQTLs | rs4844289 (chrX.70407983.A.G) |

We observed atypically long-range LD for the previously identified rare *HOXB13* rs138213197, beyond a 1Mb window from the lead variant (Supplementary Figure 5). This observation was substantiated by considerable extended haplotype homozygosity for the rare missense allele (Figure 2a). In particular, rs138213197 had an integrated haplotype score (iHS) equal to 2.87 (iHS_L_: 3.53, iHS_R_: 2.54) in our HRC haplotype data, greater than the nominal |iHS| > 2 threshold, reflecting the recent origin or selective constraint at the rs138213197 locus. Likewise, for the novel rare variant rs555778703, the rare G risk allele (Figure 2b) had an iHS equal to 2.31 (iHS_L_: 2.00, iHS_R_: 2.79). For a proxy variant rs57029021 (LD r^2^ = 0.666 in 1000 Genomes Project Phase III EUR) of the novel rare variant rs557046152 (which was unmeasured in the EGA HRC reference genomes), the rare A allele had an iHS equal to 0.87 (iHS_L_: 1.60, iHS_R_: 0.77; Figure 2c).

**Figure 2.**
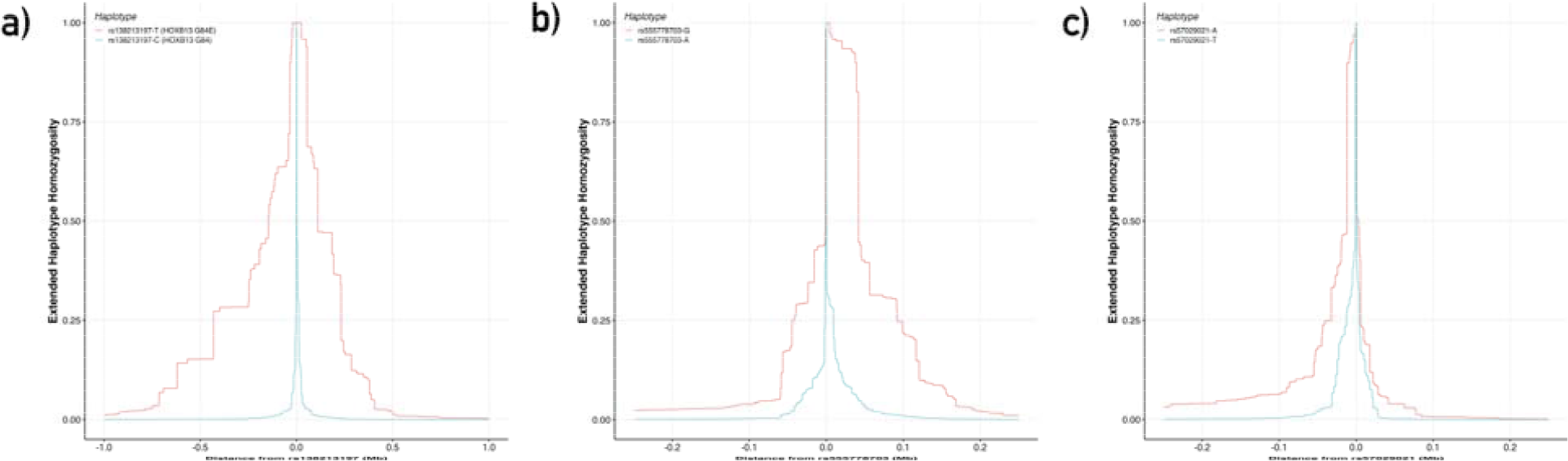
Extended Haplotype Homozygosity of Prostate Cancer Associated Rare Variants. “Haplotype Lengths for Rare PrCa Risk Variants. Extended haplotype homozygosity (EHH) plots illustrating the decay in non-recombinant linkage (Y-axis) with increasing distance along the length of the haplotypes centered at two alleles of a “core” query variant (X-axis). Differences in EHH, iHH (the area under the EHH curve), and iHS (the log-ratio between the iHH for the derived and ancestral allele) may reflect a difference in allelic age between the derived and ancestral alleles, or alternatively the selective pressure to retain a particular allele with preference to the alternative. 2a. EHH curves for the rare *HOXB13* G84E missense variant and Northern European founder mutation rs138213197, for which the iHS value of 2.87 (iHS_L_: 3.53, iHS_R_: 2.54) reflects the more recent origin of the derived G84E allele rs138213197-T. 2b. EHH curves for the novel rare variant association rs555778703, with an his value of 2.31 (iHS_L_: 2.00, iHS_R_: 2.79). 2c. EHH curves for rs57029021, an LD proxy variant for the novel rare indel association rs557046152 (LD r^2^ = 0.666 in 1000 Genomes Project Phase III EUR) with an iHS value of 0.87 (iHS_L_: 1.60, iHS_R_: 0.77).”

### Polygenic Risk Scores and Functional Interpretation

For European-ancestry subjects in KP and UKB, there was a strong association between being in the top versus bottom decile of the PRS and prostate cancer (Supplementary Figure 6, Supplementary Table 4; OR [95% CI] = 7.66 [6.78, 8.64], *P* = 1.80*10^-239^; OR_KP_EUR_ = 6.54 [5.45, 7.85], *P* = 1.32*10^-90^; OR_UKB_EUR_ = 8.63 [7.18, 10.4], *P* = 5.49*10^-117^).

To characterize the functional consequences of common variants, we examined their effects on gene expression and transcription factor binding. Among the 187 previously reported PrCa risk variants and 3 novel risk variants identified, 28 were in linkage disequilibrium (LD r^2^ > 0.5 in 1000 Genomes Project Phase III EUR) with an expression quantitative trait locus (eQTL) variant in our regularized models of normal prostatic expression levels (Table 2). Furthermore, 21 variants were predicted to significantly alter transcription factor binding site (TFBS) affinities (Table 3). rs2680708 (17q22) showed the greatest fold change in predicted binding affinity (log-difference *P*_Binding_ = 6.09) of any variant-TF pair analyzed (Table 3).

**Table 3.** Predicted Impact of PrCa Risk Variants on Transcription Factor Binding Affinity

| Previously Reported PrCa Risk Variant<br>*Genomic Locus*<br>*Risk Allele (Alt)* | Transcription Factor | $P^{\text{Binding}}$<br>Risk Allele | $P^{\text{Binding}}$<br>Alt Allele | Log-Difference in Transcription Factor Binding<br>*P-value Between Risk Allele and Alternate Allele* | TRANSFAC Vertebrate 2010.1 Matrix Name |
| --- | --- | --- | --- | --- | --- |
| rs2680708<br>17q22<br>G (A) | AR | 0.49 | 3.91E-07 | 6.09 | AR_Q6 |
|  | DBP | 0.13 | 3.91E-07 | 5.53 | DBP_Q6 |
| rs5799921<br>12q21.33<br>GA (G) | HMGIIY | 0.37 | 3.38E-07 | 6.04 | HMGIIY_Q6 |
| rs2660753<br>3p12.1<br>T (C) | AP1 | 0.55 | 2.10E-06 | 5.42 | AP1_Q6_01 |
|  |  | 0.42 | 2.10E-06 | 5.3 | AP1_Q4_01 |
| rs7210100<br>17q21.33<br>A (G) | DELTAEF1 | 0.57 | 2.31E-06 | 5.39 | DELTAEF1_01 |
| rs9600079<br>13q22.1<br>T (G) | TATA | 8.98E-07 | 0.15 | 5.22 | TATA_C |
| rs742134<br>22q13.2<br>G (A) | STAT5A | 0.01 | 4.03E-08 | 5.12 | STAT5A_03 |
|  | HNF1 | 2.70E-03 | 4.03E-08 | 4.83 | HNF1_Q6_01 |
| rs9625483<br>22q12.1<br>A (G) | MAFB | 1.56E-06 | 0.2 | 5.1 | MAFB_01 |
| rs5759167<br>22q13.2<br>G (T) | DBP | 0.07 | 9.55E-07 | 4.84 | DBP_Q6 |
| rs10086908<br>8q24.21<br>T (C) | GATA3 | 2.39E-06 | 0.07 | 4.47 | GATA3_01 |
| rs59308963<br>2q33.1<br>- (ATTCTGTC) | TCF11 | 1.44E-05 | 0.37 | 4.41 | TCF11_01 |
| rs1935581<br>10q23.31<br>C (T) | STAT1 | 3.43E-06 | 0.06 | 4.27 | STAT1_03 |
|  | STAT4 | 3.43E-06 | 0.05 | 4.16 | STAT4_01 |
| rs4245739<br>1q32.1<br>A (C) | HNF4 | 6.83E-05 | 0.98 | 4.16 | HNF4_Q6_02 |
| rs1283104<br>3q13.12<br>G (C) | FXR | 7.42E-04 | 5.35E-08 | 4.14 | FXR_Q2 |

| Previously<br>Reported<br>PrCa Risk<br>Variant<br>*Genomic Locus*<br>*Risk Allele (Alt)* | Transcription<br>Factor | $P_{\text{Binding}}$<br>Risk Allele | $P_{\text{Binding}}$<br>Alt Allele | Log-Difference<br>in Transcription<br>Factor Binding<br>*P*-value Between<br>Risk Allele and<br>Alternate Allele | TRANSFAC<br>Vertebrate 2010.1<br>Matrix Name |
| --- | --- | --- | --- | --- | --- |
| rs76551843<br>5q35.1<br>A (G) | IPF1 | 4.73E-05 | 0.58 | 4.09 | IPF1_Q6 |
| rs6869841<br>5q35.2<br>A (G) | HOXA3 | 0.29 | 4.25E-05 | 3.84 | HOXA3_01 |
| rs13385191<br>2p24.1<br>G (A) | NFAT1 | 0.39 | 5.95E-05 | 3.82 | NFAT1_Q6 |
| rs1571801<br>9q33.2<br>A (C) | ZNF333 | 4.16E-05 | 0.27 | 3.82 | ZNF333_01 |
| rs339331<br>6q22.1<br>T (C) | IRF8 | 0.37 | 6.58E-05 | 3.75 | IRF8_Q6 |
| rs1283104<br>3q13.12<br>G (C) | PNR | 2.66E-04 | 5.35E-08 | 3.7 | PNR_01 |
| rs182314334<br>3q25.1<br>T (C) | POU1F1 | 0.04 | 7.81E-06 | 3.69 | POU1F1_Q6 |
| rs17694493<br>9p21.3<br>G (C) | STAT | 0.34 | 8.04E-05 | 3.62 | STAT_Q6 |

## DISCUSSION

We combined imputed genotype data from two large cohorts totaling 14,113 PrCa cases and 201,722 controls, with a reference panel of over 27,000 phased genomes, to investigate the effects of rare genetic variants and the mechanisms and cumulative impact of common variants on PrCa risk. Three novel loci, including two rare variants (rs557046152 at 8p12, rs555778703 at 4q31.21) and one common variant (rs62262671 at 3p21.31), were associated with PrCa in our meta-analysis of European-ancestry subjects across cohorts. Likewise, an additional three novel variants were associated with PrCa in our meta-analysis, although this finding was driven primarily by the UKB participants.

Furthermore, the PRS associations we observed for European ancestry men were of larger magnitude of effect than reported previously, when there were only 105 known PrCa risk variants [6]. Namely, the nearly 8-fold increase in PrCa risk for men in the top vs. lowest decile of the PRS suggests that such a score may have similar predictive ability as high penetrance genes used to predict cancer risk in clinical practice, such as *BRCA1* (OR [95% CI]: 5.91 [5.25, 6.67]) and *BRCA2* (OR [95% CI]: 3.31 [2.95, 3.71]) for breast cancer risk [27]. Although the PRS effect is of relatively large magnitude, the scores may not be transferable to subjects of non-European descent [6], can be biased by genetic drift between ethnic groups [28], and could potentially widen existing health disparities [29].

Integration of gene expression and transcription factor binding site affinity data suggested novel mechanisms for many of the common PrCa variants previously reported. One example is a highly significant change in binding affinity at rs2680708. This finding is especially interesting given that rs2680708 abrogates a binding site for Androgen Receptor, a master regulator of prostatic gene expression. While our functional analyses did not nominate any genes whose expression may be affected as a consequence of eliminating this particular binding site, further study may reveal the effect of rs2680708 on the dysregulation gene expression or additional molecular processes. We also identified a putative mechanism of Oct1 binding for the newly implicated rs62262671 risk variant (3p21), which was predicted to have a large impact (log-difference *P*_Binding_ = 3.03) on binding affinity for Oct1, a TF with a known impact on PrCa and Androgen Receptor signaling [30]. Given that rs62262671 was also identified as an eQTL affecting the expression of *RBM6* and *UBA7*, these findings suggest that Oct1 may be involved in the regulation of the expression of these two genes, and provides a hypothesis for future functional follow-up regarding the involvement of these genes in prostate cancer development.

The mechanisms through which the rare, noncoding variants we identified are associated with prostate cancer remain somewhat unclear, with a lack of precise functional evidence regarding mechanism of action or close proximity to genes or known risk loci in *cis*. This underscores the challenge of not only detecting—but also interpreting—how rare variants impact the genetic etiology of complex traits using existing gene-based methodology and functional genomic datasets. Improved functional datasets may clarify the effects of rare variants on expression, splicing, or methylation.

Selective scans, which use population genetics metrics (such as EHH and iHS) to identify signatures of positive or negative natural selection [21], face similar challenges—rare variants naturally reside on longer haplotypes and obscure the direction of any selective forces that may act upon them [31]. If polymorphisms more exclusive, or even private, to a particular lineage or family comprise a substantial portion of disease risk for PrCa (or other traits), then new approaches and assays for both detecting and characterizing the relevant anomalies of these causal variants will be needed. These considerations are of particular importance given the proliferation of rare polymorphisms as a result of recent explosive human population expansion [32]. Hence, with the majority of all human variation shifting towards the low end of the allele frequency spectrum, identifying operative aberrations poses a significant challenge.

In spite of these challenges, over a decade of GWAS efforts [33] has advanced the genetic characterization of prostate cancer considerably. Our implementation of a PRS model for PrCa demonstrates this remarkable progress and the predictive power of aggregating PrCa risk loci.

## CONCLUSIONS

By undertaking a GWAS in the large KP and UKB population-based cohorts, we detected multiple novel PrCa risk loci, including two rare variants, rs557046152 and rs555778703. Our PRS analysis of known common PrCa risk variants indicated that European ancestry men in the highest PRS decile have a substantially increased risk that may be of clinical importance; however, the result was greatly attenuated in other ancestries. Functional characterization of PrCa risk variants using gene expression and transcription factor binding affinity data revealed putative mechanisms. However, further study is needed to more fully illuminate the biological interactions that facilitate the influence of PrCa risk loci, in particular for rare variants.

## AUTHOR CONTRIBUTIONS

John S. Witte had full access to all the data in the study and takes responsibility for the integrity of the data and the accuracy of the data analysis.

*Study concept and design:* Emami, Risch, Hoffmann, Van Den Eeden, Witte.

*Acquisition of data:* Cario, Mefford, Wan, Wong, Aaronson, Presti, Habel, Shan, Ranatunga, Chao, Ghai, Jorgenson, Sakoda, Kwok, Schaefer, Risch, Van Den Eeden, Witte.

*Analysis and interpretation of data:* Emami, Cavazos, Rashkin, Cario, Tai, Mefford, Kachuri, Kvale, Hoffmann, Witte.

*Drafting of the manuscript:* Emami, Graff, Witte.

*Critical revision of the manuscript for important intellectual content:* Emami, Hoffmann, Witte. *Statistical analysis:* Emami, Cavazos, Rashkin, Witte.

*Obtaining funding:* Schaefer, Risch, Van Den Eeden, Witte. Administrative, technical, or material support: None. *Supervision:* Witte.

*Other (specify):* None.

*Financial disclosures:* John S. Witte certifies that all conflicts of interest, including specific financial interests and relationships and affiliations relevant to the subject matter or materials discussed in the manuscript (e.g. employment/affiliation, grants or funding, consultancies, honoraria, stock ownership or options, expert testimony, royalties, or patents filed, received, or pending), are the following: None.

*Funding/Support and role of the sponsor:* Supported by the Robert Wood Johnson Foundation, National Institutes of Health (R01CA088164, R01CA201358, R25CA112355, T32GM067547, and U01CA127298), the UCSF Goldberg-Benioff Program in Cancer Translational Biology, the UCSF Discovery Fellows program, the Microsoft Azure for Research program, and the Amazon AWS Cloud Credits for Research program. Support for participant enrollment, survey completion, and biospecimen collection for the RPGEH was provided by the Robert Wood Johnson Foundation, the Wayne and Gladys Valley Foundation, the Ellison Medical Foundation, and Kaiser Permanente national and regional community benefit programs. Genotyping of the GERA cohort was funded by a grant from the National Institute on Aging, the National Institute of Mental Health, and the NIH Common Fund (RC2 AG036607). The sponsors played no role in the study.

## Acknowledgments

The authors thank Elad Ziv, Nadav Ahituv, and Ryan Hernandez for their guidance and feedback. Furthermore, we are grateful to the Kaiser Permanente Northern California members who have generously agreed to participate in the Kaiser Permanente Research Program on Genes, Environment, and Health, the ProHealth Study and the California Men’s Health Study. This research has been conducted using the UK Biobank Resource under Application Number 14105.

## APPENDIX A.

Supplementary data

Supplementary data associated with this article can be found at the journal online.

## SUPPLEMENTARY MATERIALS

### Custom Microarray Design and Genotyping

In our design of a DNA microarray with predominantly custom, functionally relevant markers, the SNP selection procedure was conducted as follows. First, a set of target markers was constructed. This target set included variants previously associated in genome-wide association studies (GWAS), significant and suggestive, of prostate cancer (PrCa) associated traits (PSA level, gene-by-gene interactions), other correlated traits (breast cancer, height, body mass index, obesity, diabetes, and others), and also uncorrelated traits (all NHGRI GWAS catalog traits). Additionally, a set of pan-cancer candidate genes was compiled by experimental colleagues, and all rare variants in windows centered around these genes were included in the target set. Rare variant in windows around highly mutated genes from the somatic cancer database COSMIC were also included. Furthermore, rare variants from a series of whole genome and whole exome sequence analyses (of African American PrCa case normal genomes [1], The Cancer Genome Atlas (TCGA) [2] and dbGaP [3] normal exomes, and ENCODE PrCa DNAse I hypersensitive regions) were put into the target set.

Second, variant selection was conducted with complementarity to the GWAS array previously assayed in the study population in order to limit redundancy (Supplementary Figure 1b), drawing from a candidate set disjoint from the GWAS array markers. This produced a set of primarily rare selected markers optimized for coverage of the target set, through tagging and direct genotyping (Supplementary Figure 1a; Supplementary Table 2).

Genotyping sample DNA plates without special attention to matching case and control covariates can lead to batch effects. In order to minimize batch effects and expedite genotyping, a sample management system (Samasy) [4] and sample selection algorithm were designed and implemented to robot-automate DNA sample allocation. The greedy sample selection algorithm for moving case and control DNA samples from source plates to destination plates was designed with the following objectives: 1) Use all available PrCa cases on source plates, 2) Select equal numbers of controls and cases, 3) Frequency match the distribution of race and age in controls and cases, while oversampling African American controls and rare (race and age) strata to improve power, 4) Select all required samples from a source plate at one time so lab workers will only have to locate and handle a source plate once, 5) Optimize work flow so sets of source and destination plates can be simultaneously loaded and unloaded from the Biomek liquid handling robot.

### Quality Control and Imputation

The quality control (QC) process is described in Supplementary Figures 2a and 2b.

In order to produce the highest confidence genotype calls for the greatest number of samples and probesets, sample quality was first evaluated to screen for and eliminate potential outliers that may negatively impact downstream genotype clustering. Sample QC was executed in three stages. First, signal-to-noise (DishQC (DQC), ranging from 0 to 1) was computed for each sample based on intensity data calculated from the raw microarray fluorescence images. A threshold was drawn to exclude samples with low signal-to-noise (DQC < 0.75) based on the inflection point of the empirical distribution of DQC values. Second, 20,000 diagnostic Step 1 probesets were genotyped using 5 sample batches (grouped chronologically based on sample processing dates) to obtain empirical distributions of sample call rates for each batch. A call rate threshold was drawn based on the inflection points of these distributions to exclude samples with low call rates (CR < 0.95) or missing covariates from further analyses, leaving 14,818 samples. Third, the remaining samples were re-genotyped at all 416,047 Step 2 probesets using the same chronological packaging. Correlations between heterozygosity rate and call rate were used to define three clustered tiers (Supplementary Figure 2a) of sample quality: higher quality samples (HQ, with Call Rate > 97% and Heterozygosity Rate < 15%), lower quality samples (LQ), and plate quality control (PQC) samples (Plate Call Rate < 96.5%). Finally, among 80 pairs of samples with a high, 1st degree level of relatedness (kinship coefficient > 0.35), one of each pair of individuals was removed from further analysis, preferentially retaining prostate cancer cases. This concluded sample QC and provided a basis for evaluating QC of probesets through additional re-genotyping.

Given the remaining 14,818 samples, each labeled according to three tiers of sample quality outlined above, re-genotyping was performed to separate well resolved probesets from those susceptible to batch effects, and strategies were implemented to correct for these batch effects for the greatest number of probesets. First, all 14,818 samples, regardless of tier, were genotyped across all 416,047 probesets. Genotype clusters were next evaluated across all probesets, and classified as being either well resolved across all samples (i.e. not susceptible to batch effects, n=327,703), well resolved across only HQ samples (n=6,672), or poorly resolved (n=81,672). In order to remedy batch effects, first the probesets resolved in the HQ samples were considered. An Empirical Bayes genotyping strategy was implemented in which the well resolved HQ genotype clusters were used to sequentially guide the genotyping of samples in the lower quality LQ and PQC tiers. By packaging LQ tiers with HQ tiers, and using the posterior HQ cluster centers as AxiomGT1 [5] priors for re-genotyping, genotype calls were produced for each probeset across all sample tiers. Minor allele frequency (MAF) was compared among sample tiers in order to identify probesets where call frequencies were in agreement and in disagreement. Genotype calls for which MAF agreed among tiers (within 10% of HQ samples MAF) were retained as final genotypes for their respective probesets, while those probesets exhibiting MAF disagreements among tiers were combined with the other poorly resolved probesets for a series of procedures.

These procedures included re-thresholding the genotype cluster metric Heterozygous Strength Offset, which measures the displacement of the heterozygous cluster in relation to the homozygous clusters, as well as using linear regression to normalize probeset intensities across plates. The latter procedure led to the reclassification of 15,319 probesets as well resolved, and 65,177 as poorly resolved. These poorly resolved variants were processed through additional steps (Supplementary Figure 2b) to identify monomorphic probesets based on a genotype cluster metric Homozygote Ratio Offset (HomRO) and isolate calls for only HQ samples. The probesets well resolved in only HQ samples, in addition to those reclassified by plate-normalization, were combined with the remaining well-resolved probesets for optimization of both polymorphic probeset detection sensitivity (Minor Homozygote, Het Cluster Strengthening) and also rare variant detection sensitivity (Rare Variant Per-Plate Re-Genotyping).

After the conclusion of the preceding raw genotype QC steps described above and outlined in Supplementary Figure 2b, several additional QC steps were performed on the called genotypes for the genotype-resolved variants (n=356,671). First, variants that deviated from Hardy-Weinberg Equilibrium (*p* < 1e-4) in European ancestry controls were removed (n=30,632), leaving n=326,039 variants. Next, to further control for batch effects, variants where genotype was associated with sample quality tier (HQ vs. LQ; *p* < 1e-4) were further excluded (n=1,376). Furthermore, variants where European ancestry minor allele frequency deviated from the Thousand Genomes Project EUR MAF [6] by greater than 10% (n=164) or that were monomorphic (n=69,431) were filtered out, leaving n=255,068 variants remaining. Furthermore, variants with low AxiomGT1 genotype clustering confidence scores (less than 0.2) were identified (n=22,560) and excluded from further analysis. Finally, variants which evaded prior QC but whose cluster plots upon visual inspection raised suspicion of a cluster split (i.e. misclustering by the AxiomGT1 algorithm, leading to misclassification of homozygotes as heterozygotes, or vice versa) were excluded (n=100), yielding a final total of 232,408 variants for phasing, imputation, and downstream analysis.

### Evolutionary History of Rare Variants

For generating a genetic map to be used in calculating EHH and iHS, the predictGMAP program [7] was used to interpolate genetic map positions, using 1000 Genomes Project OMNI genetic map files as a reference [8]. The selscan package [9] was run with the settings “--keep-low-freq”, “--max-extend 0”, “--threads 8”, and “--ehh-win 500000,” with the exception that “--ehh-win 1000000” was invoked for the HOXB13 G84E mutation rs138213197 to account for longer range LD. The integrative haplotype score (iHS) was computed manually using a python script implementing equations (4) and (5) from the selscan publication [9].

### Polygenic Risk Score Analyses

Polygenic risk scores (PRS) of PrCa risk were computed by taking sum of the effect sizes for 187 previously reported PrCa risk loci [10-31] (Table 3). This included the 105 variants previously modeled by Hoffmann et al. in 2015 [32], the 63 novel variants discovered by Schumacher et al. in 2018 in the European-ancestry PRACTICAL consortium [33], as well as summary statistics reported for an additional 20 independent variants (LD r^2^≤ 0.3 in 1000 Genomes Project Phase III EUR) [34-38].

### Functional Annotation

To interpret the functional relevance of the known PrCa risk variants, we trained elastic net regression models of normal prostatic gene expression [39]. Training samples with paired genotype and gene expression data were drawn from the National Center for Biotechnology Information (NCBI) publicly available database of Genotypes and Phenotypes (dbGaP phs000985.v1.p1). Training data derived from a study which collected histologically normal prostate tissue from consenting subjects (471 European-ancestry men; mean age [SD]: 60.1 [7.15] for the 249 men with age available) having undergone radical prostatectomy treatment for prostate cancer (N = 453; 63.6% Gleason 6, 36.4% Gleason 7) or cystoprostatectomy treatment for bladder cancer (N = 18). Inclusion criteria, histopathological assessment, sample processing, and quality control were described previously for these data [40].

We imputed unobserved training data genotypes to the 1000 Genomes Project Phase III reference panel using a pre-phasing workflow to match the strand and reference allele recorded in the data with those observed in the reference panel, while excluding ambiguous variants and indel mutations. Next, samples were phased and imputed using Eagle v2.3 [41] (cohort-based) and Beagle v4.1 [42], respectively. Gene boundaries (hg38) for 17,233 transcripts were downloaded from the NCBI Gene database using the Biopython Entrez eutils REST API [43]. Genomic coordinates were converted from hg38 to hg19 (GRCh37) via UCSC liftOver. For each transcript, well-imputed (r^2^_INFO_ > 0.8) training data genetic variants located (a) in the within 500kb of the start position, (b) between the start and end positions, inclusive, or (c) within 500kb of the end position, were extracted. Next, following the PrediXcan gene expression model training procedure [44], a regularized regression model was fit with the R (v3.2.2) package GLMNet [45], with genetic variants *in cis* to a given transcript as the design matrix, and the transcript RNA-Seq RPKM levels as the response variable. Models with at minimum one non-intercept explanatory variable retained were produced for 13,258 genes, and leave-one-out cross validation (LOOCV) was utilized (loss function: R cv.glmnet type.measure = “mse”) to select coefficients minimizing mean cross-validated error (regularization parameter: R predict s = “lambda.min”).

To examine allele-specific effects on transcription factor binding site affinity for the set of known PrCa variants, 25 base pair 3’ and 5’ flanking sequences were downloaded from the UCSC table browser [46] via Selenium webdriver automation. Next, FASTA sequences containing both the major and minor variant alleles were automatically analyzed through the sTRAP Transcription Factor Affinity Prediction webserver [47], with parameters “matrix file” = “transfac_2010.1 vertebrates”, “background model” = “human_promoters”, and “Multiple test correction” = “Benjamini-Hochberg.”

---

**Supplementary Table 1.** Description of the Study Cohort

| Ethnic Group |  | Kaiser Permanente |  | UK Biobank |  |
| --- | --- | --- | --- | --- | --- |
|  |  | Cases | Controls | Cases | Controls |
| European Ancestry | N | 6,196 | 5,453 | 7,917 | 188,352 |
|  | Age [SD] | 68.1 [7.9] | 71.5 [10.8] | 64.1 [5.6] | 57.1 [8.1] |
|  | BMI [SD] | 26.9 [4.2] | 27.0 [4.4] | 27.6 [4.0] | 27.8 [4.6] |

\* Subjects restricted to unrelated individuals.

**Supplementary Table 2.** Custom Microarray Design Modules

| Module Name | Number of Variants* | Module Description |
| --- | --- | --- |
| Missense | 67,846 | Nonsynonymous coding mutations |
| Tag | 57,607 | Variants tagging targeted content in other modules |
| Bad (EUR) | 33,646 | GWAS markers that did not previously pass QC on Affy EUR Array [1] |
| Bad (AFR) | 33,584 | GWAS markers that did not previously pass QC on Affy AFR Array [2] |
| Fine Mapping | 29,096 | Variants selected from previously reported prostate cancer GWAS loci |
| LOF | 24,783 | Loss-of-function coding mutations |
| WES_TCGA | 24,167 | Rare variants from TCGA prostate cancer patient normal tissue exomes [3] |
| Witte_somatic | 17,792 | Rare variants in windows around genes from important cancer pathways |
| CandGene4 | 17,095 | Rare variants in windows around cancer-related candidate genes (4<sup>th</sup> tier) |
| Exome319_tier2 | 16,366 | Variants from the Affy Exome319 exome chip (2<sup>nd</sup> tier) |
| CandGene1 | 13,586 | Rare variants in windows around cancer-related candidate genes (1<sup>st</sup> tier) |
| CandGene3 | 9,465 | Rare variants in windows around cancer-related candidate genes (3<sup>rd</sup> tier) |
| A_A_Rare | 9,394 | Rare variants from African American prostate cancer patient normal exomes |
| HSS | 9,081 | Rare variants from ENCODE PrCa cell line DNase I hypersensitive regions |
| HGMD | 8,557 | Rare variants from HGMD database gene regions |
| Cosmic | 7,233 | Rare variants from recurrently somatically mutated cancer genes |
| CandGene5 | 7,095 | Rare variants in windows around cancer-related candidate genes (5<sup>th</sup> tier) |
| tier2 eQTL | 6,918 | Variants associated with gene expression levels |
| cancer | 6,706 | UK Biobank cancer variation module |
| Exome319_tier1 | 6,402 | Variants from the Affy Exome319 exome chip (1<sup>st</sup> tier) |
| Neanderthal | 5,680 | Variants thought to be introduced during Human-Neanderthal introgression |
| LOF novel | 5,194 | Loss-of-function coding mutations |
| HGMD novelprescue | 5,132 | Rare variants from HGMD database gene regions |
| 8k eQTL | 4,662 | Variants associated with gene expression levels in different tissues |
| GWAS compatibility | 3,992 | Variants for boosting GWAS coverage |
| CandGene2 | 2,597 | Rare variants in windows around cancer-related candidate genes (2<sup>nd</sup> tier) |
| WES_dbgap | 1,426 | Rare variants from dbGaP prostate cancer patient normal tissue exomes [4] |
| HLA/KIR | 1,408 | Variants in HLA / KIR genes |
| ADME | 1,183 | Pharmacogenomics variants |
| chrY | 806 | Variants located on chromosome Y |
| ASHG | 742 | Variants from cancer genes presented at 2013 ASHG conference |
| AfAmrImputed | 583 | African American GWAS imputed variants |
| GWAS_enrichment_tier1.4 | 454 | Variants from the NHGRI GWAS catalog (4<sup>th</sup> tier) |
| KIR | 418 | Variants in the KIR gene |
| BioBank1_LoF_tier1 | 367 | Loss-of-function coding mutations |
| Kaiser GWAS | 283 | Variants associated in GWAS in the Kaiser Permanente RPGEH cohort |
| Diabetes_Metabohip | 274 | Variants related to metabolic, cardiovascular, and anthropometric traits |
| Telomere | 261 | Variants associated with leukocyte telomere length |
| GWAS_enrichment_tier1.1 | 243 | Variants from the NHGRI GWAS catalog (1<sup>st</sup> tier) |
| Height | 221 | Variants associated with height |
| Diabetes_GWAS | 211 | Variants associated with diabetes |
| ApoE | 203 | Variants in the ApoE gene |
| BrCa | 183 | Variants associated with breast cancer risk |
| chrMT | 180 | Variants located on the mitochondrial chromosome |
| GWAS_enrichment_tier1.3 | 140 | Variants from the NHGRI GWAS catalog (3<sup>rd</sup> tier) |
| Ovarian | 137 | Variants associated with ovarian cancer risk |
| Blood eQTL | 125 | Variants associated with gene expression levels in blood cells |
| GWAS_enrichment_tier1.2 | 51 | Variants from the NHGRI GWAS catalog (2<sup>nd</sup> tier) |
| CandGene0 | 50 | Rare variants in windows around cancer-related candidate genes (0<sup>th</sup> tier) |
| CandGene6 | 24 | Rare variants in windows around cancer-related candidate genes (6<sup>th</sup> tier) |
| Radiogen | 4 | Variants associated with radiogenomic phenotypes |
| special | 2 | Variants to be forced on the array |
| Alzheimers | 2 | Variants associated with Alzheimer's risk |
| CandGene1 | 1 | Variants from 1<sup>st</sup> list of cancer-related candidate genes |

\* Total sums to greater than the number of genotyped probesets because certain probesets were members of multiple modules.

**Supplementary Table 3.**
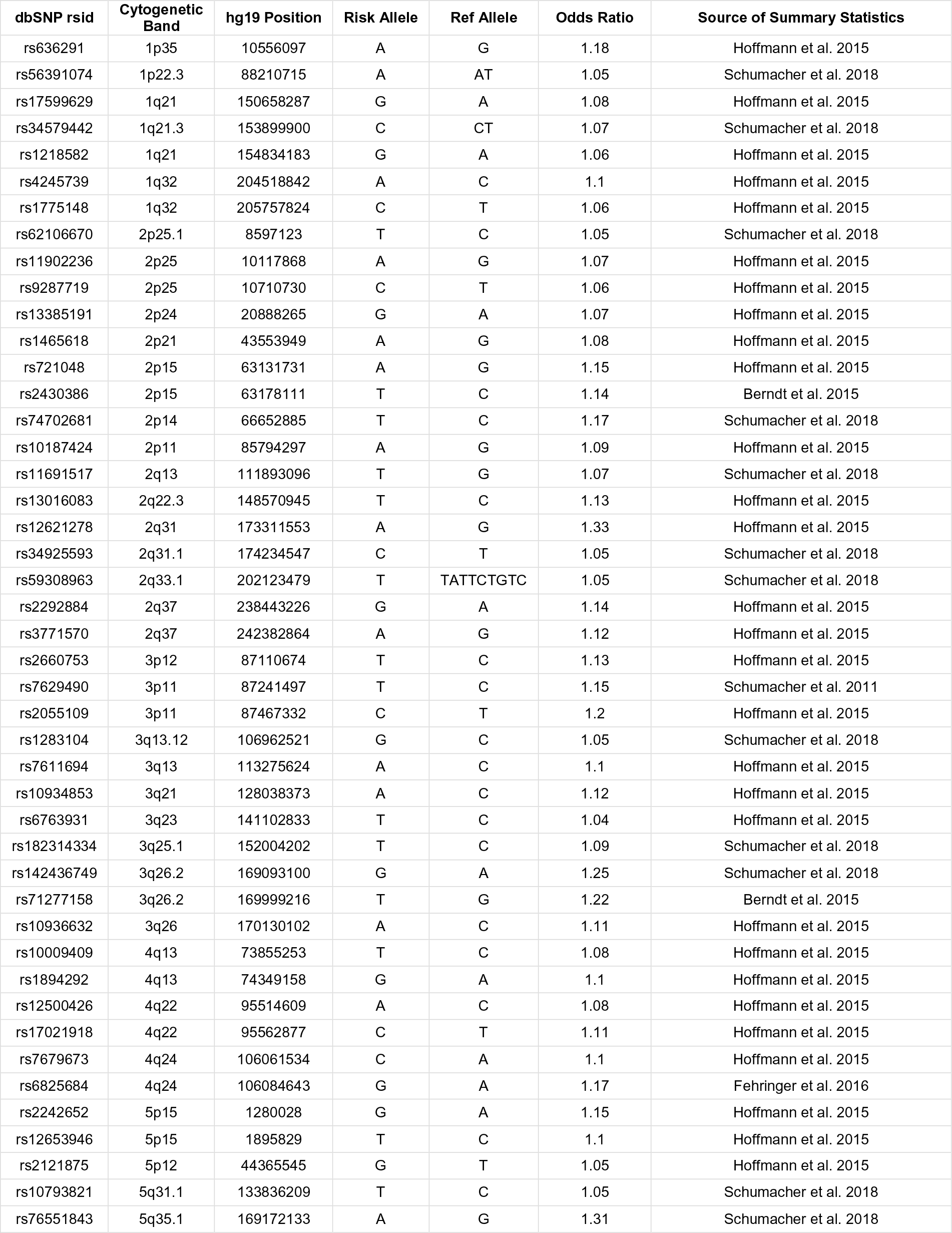
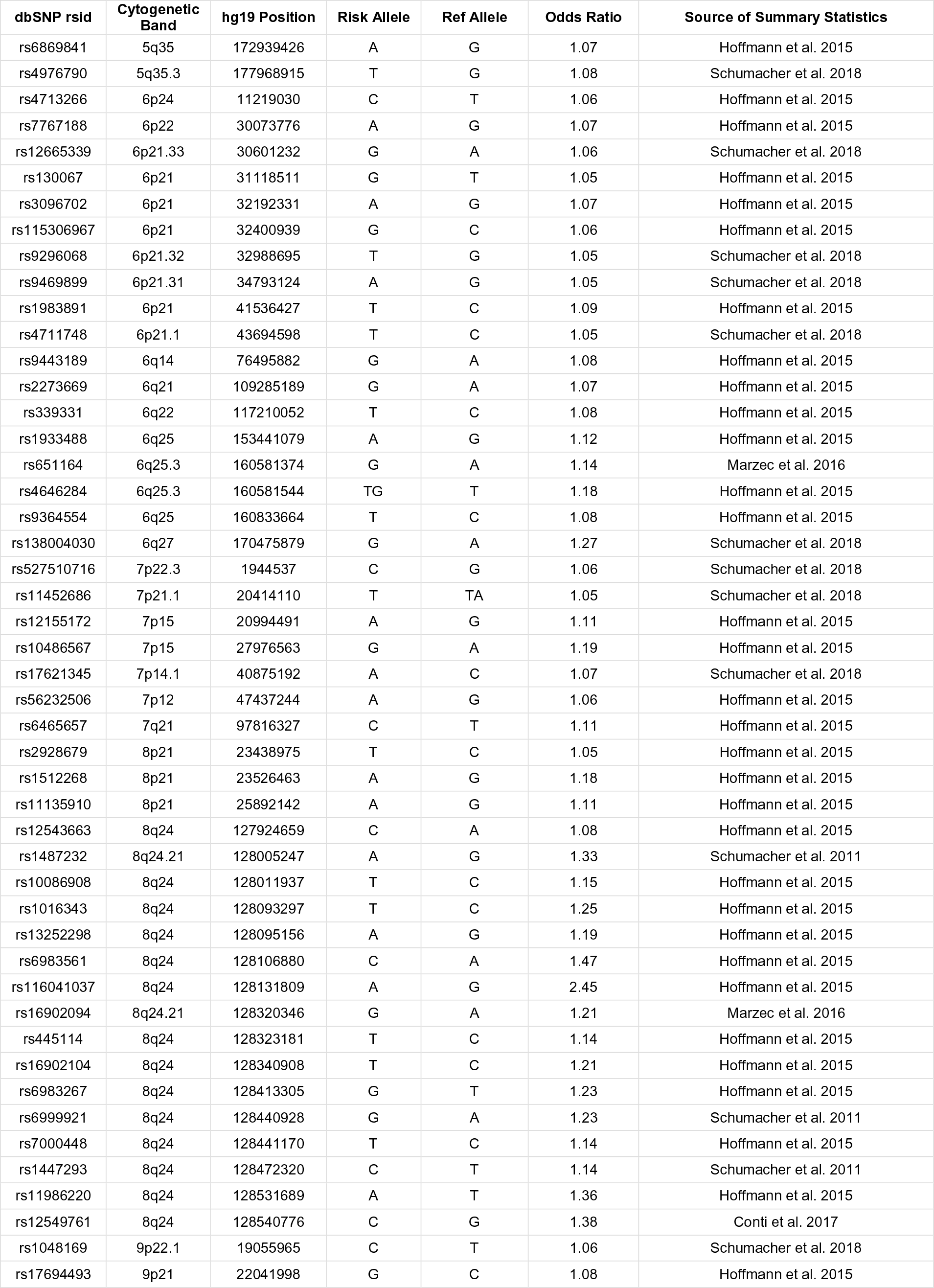
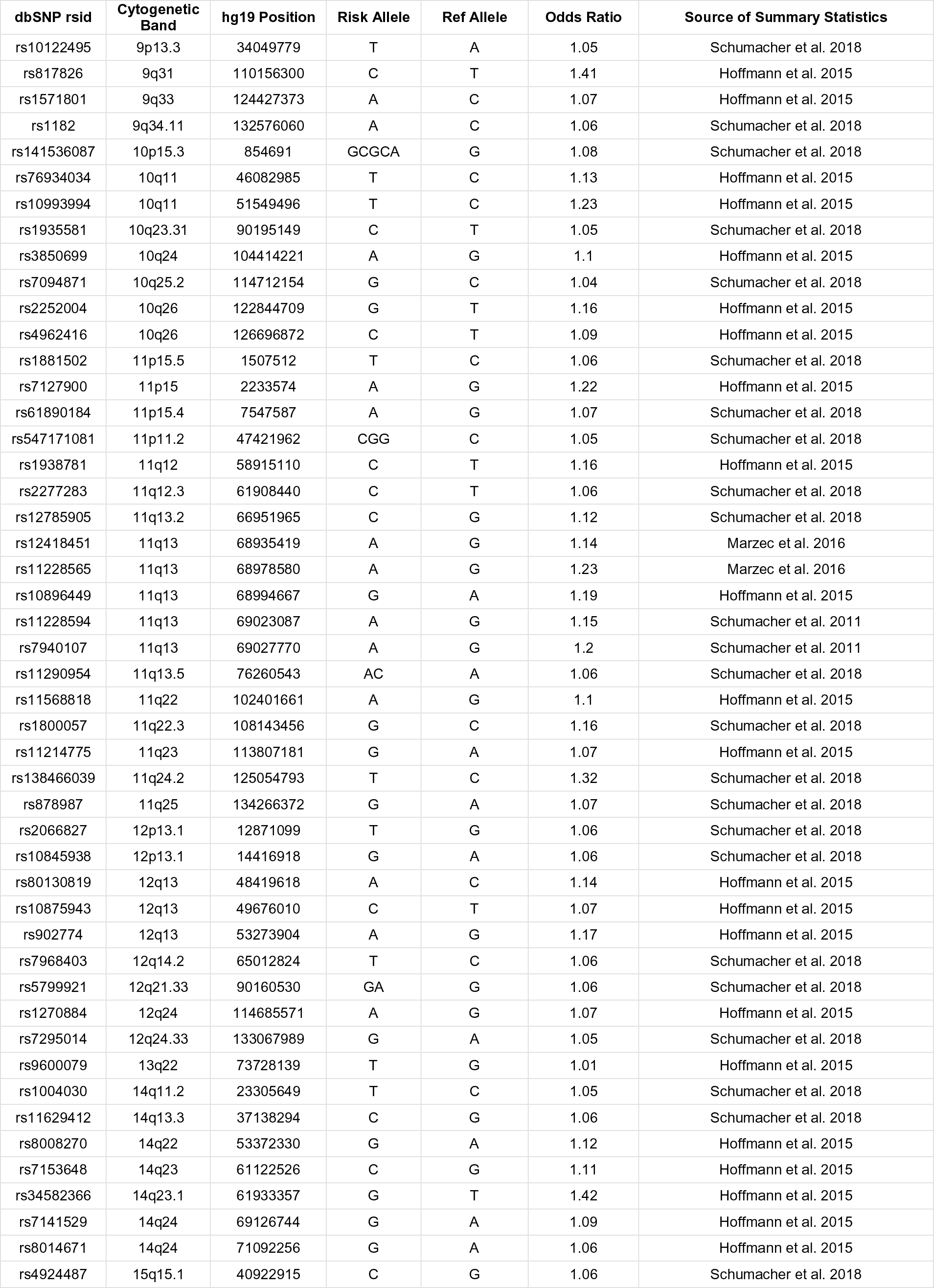
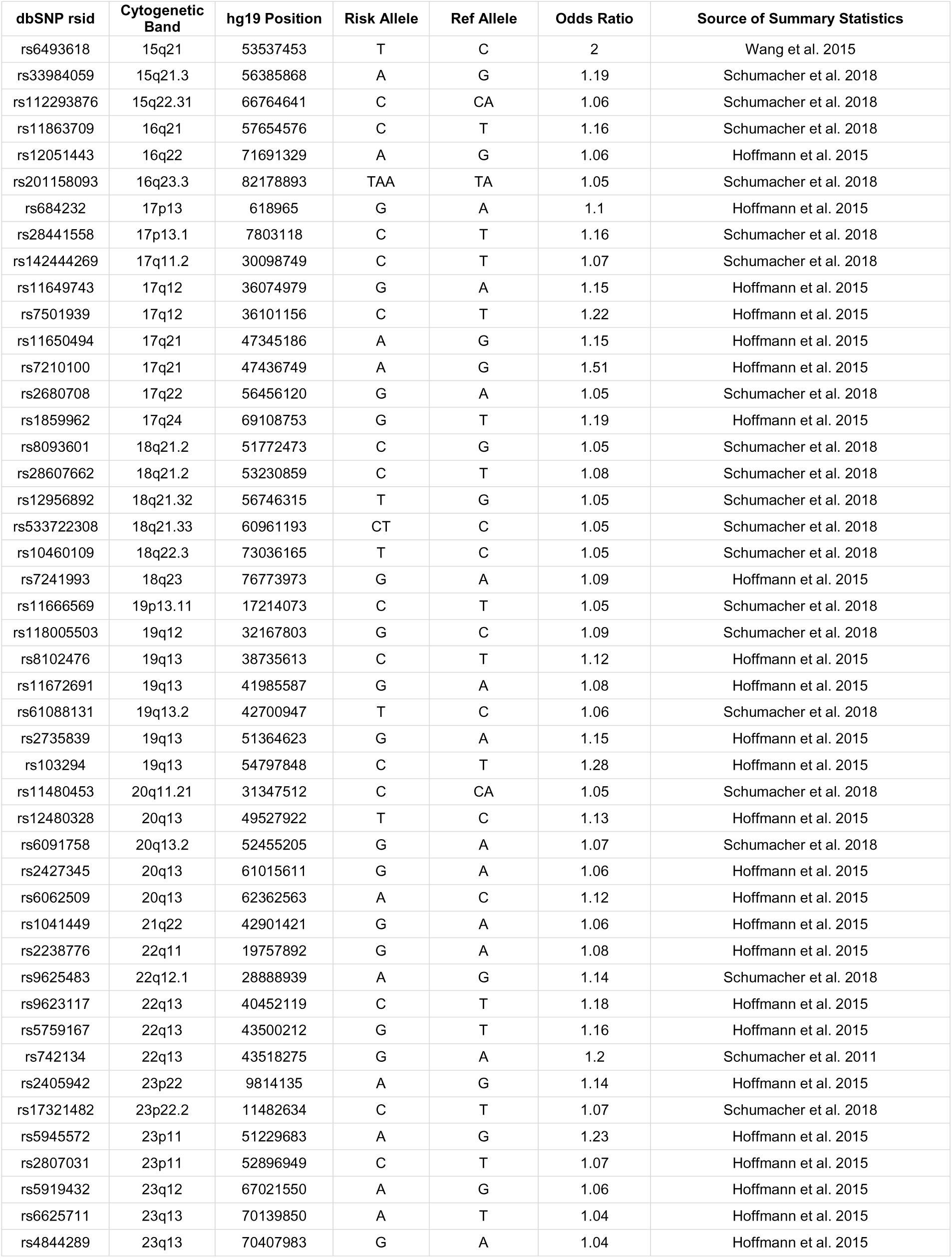
Information and Summary Statistics for 187 Variants Modeled in Prostate Cancer Polygenic Risk Score

**Supplementary Table 4.**
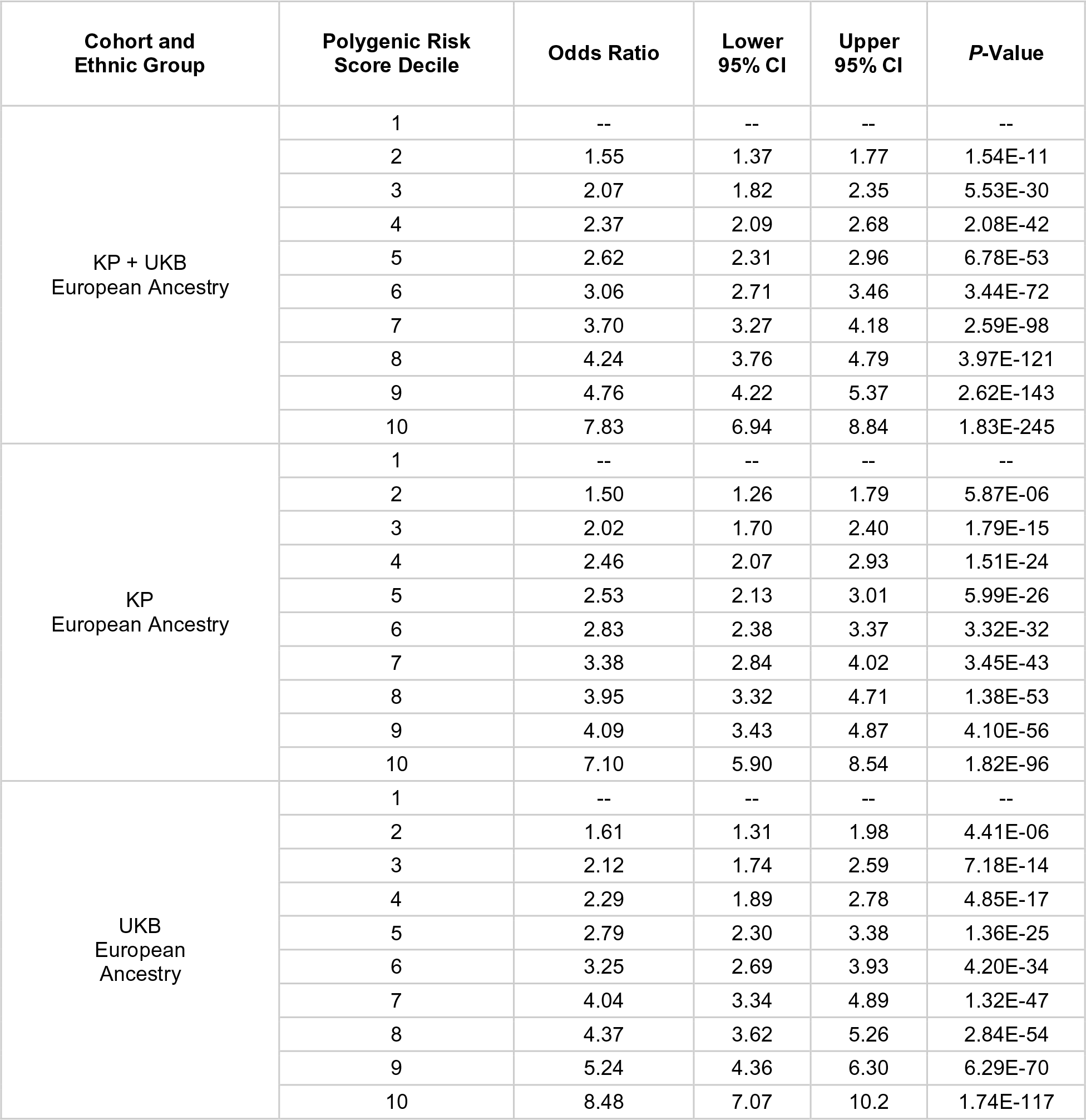
Polygenic Risk Score Performance

| Cohort and Ethnic Group | Polygenic Risk Score Decile | Odds Ratio | Lower 95% CI | Upper 95% CI | P-Value |
| --- | --- | --- | --- | --- | --- |
| KP + UKB<br>European Ancestry | 1 | -- | -- | -- | -- |
|  | 2 | 1.55 | 1.37 | 1.77 | 1.54E-11 |
|  | 3 | 2.07 | 1.82 | 2.35 | 5.53E-30 |
|  | 4 | 2.37 | 2.09 | 2.68 | 2.08E-42 |
|  | 5 | 2.62 | 2.31 | 2.96 | 6.78E-53 |
|  | 6 | 3.06 | 2.71 | 3.46 | 3.44E-72 |
|  | 7 | 3.70 | 3.27 | 4.18 | 2.59E-98 |
|  | 8 | 4.24 | 3.76 | 4.79 | 3.97E-121 |
|  | 9 | 4.76 | 4.22 | 5.37 | 2.62E-143 |
|  | 10 | 7.83 | 6.94 | 8.84 | 1.83E-245 |
| KP<br>European Ancestry | 1 | -- | -- | -- | -- |
|  | 2 | 1.50 | 1.26 | 1.79 | 5.87E-06 |
|  | 3 | 2.02 | 1.70 | 2.40 | 1.79E-15 |
|  | 4 | 2.46 | 2.07 | 2.93 | 1.51E-24 |
|  | 5 | 2.53 | 2.13 | 3.01 | 5.99E-26 |
|  | 6 | 2.83 | 2.38 | 3.37 | 3.32E-32 |
|  | 7 | 3.38 | 2.84 | 4.02 | 3.45E-43 |
|  | 8 | 3.95 | 3.32 | 4.71 | 1.38E-53 |
|  | 9 | 4.09 | 3.43 | 4.87 | 4.10E-56 |
|  | 10 | 7.10 | 5.90 | 8.54 | 1.82E-96 |
| UKB<br>European Ancestry | 1 | -- | -- | -- | -- |
|  | 2 | 1.61 | 1.31 | 1.98 | 4.41E-06 |
|  | 3 | 2.12 | 1.74 | 2.59 | 7.18E-14 |
|  | 4 | 2.29 | 1.89 | 2.78 | 4.85E-17 |
|  | 5 | 2.79 | 2.30 | 3.38 | 1.36E-25 |
|  | 6 | 3.25 | 2.69 | 3.93 | 4.20E-34 |
|  | 7 | 4.04 | 3.34 | 4.89 | 1.32E-47 |
|  | 8 | 4.37 | 3.62 | 5.26 | 2.84E-54 |
|  | 9 | 5.24 | 4.36 | 6.30 | 6.29E-70 |
|  | 10 | 8.48 | 7.07 | 10.2 | 1.74E-117 |

**Supplementary Figure 1a.**
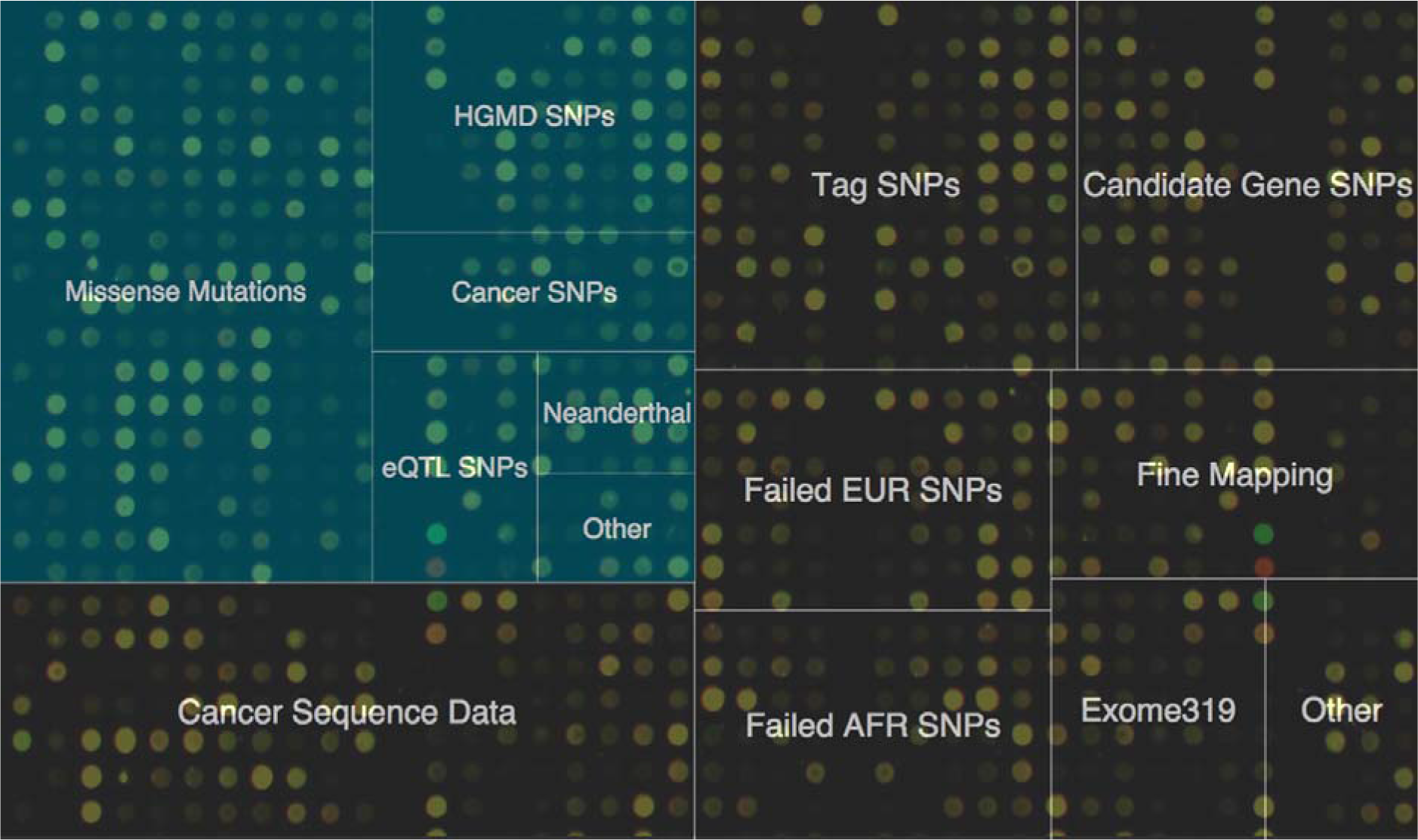
Custom Microarray Marker Content. Custom Array Design. S1a. The relative fractions of Selected Markers are grouped by their source and illustrated to scale (total of 416,047 probesets). Teal colored cells derive from the UK Biobank Array modules and include a diverse set of curated and functionally relevant mutations. S1b. SNP Selection was conducted according to a greedy algorithm. In a single iteration of the algorithm, Candidate Markers are ranked for coverage of Target Markers, the best candidate is moved to the set of Selected Markers, and the candidates are re-ranked. The algorithm allows for markers to be “pre-selected” by placement in the Selected Markers set upon initialization, and runs until the Selected Markers bin equals a certain maximum value. Probesets for the resulting selected markers are included in the fabrication of the custom microarray chip.

**Supplementary Figure 1b.**
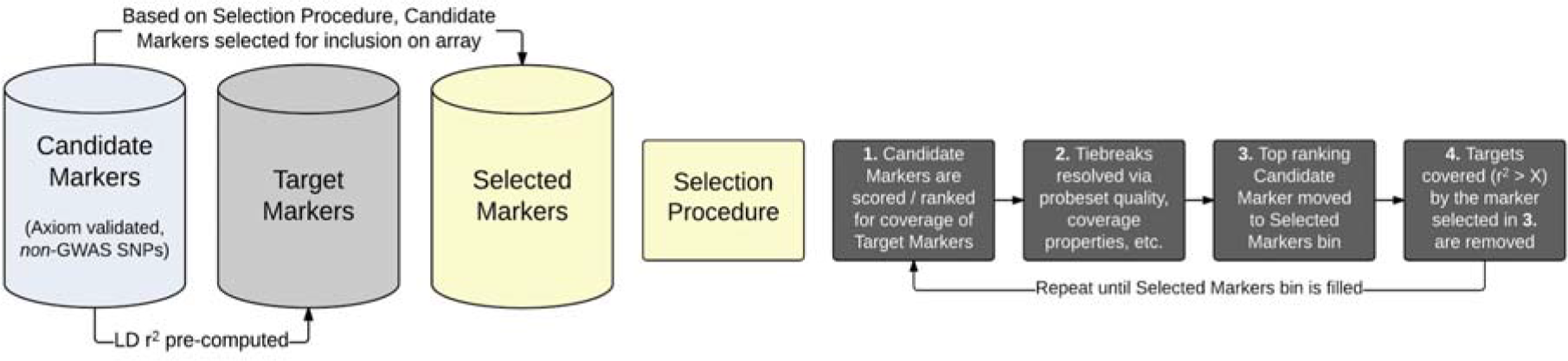
SNP Selection Algorithm. Custom Array Design. S1a. The relative fractions of Selected Markers are grouped by their source and illustrated to scale (total of 416,047 probesets). Teal colored cells derive from the UK Biobank Array modules and include a diverse set of curated and functionally relevant mutations. S1b. SNP Selection was conducted according to a greedy algorithm. In a single iteration of the algorithm, Candidate Markers are ranked for coverage of Target Markers, the best candidate is moved to the set of Selected Markers, and the candidates are re-ranked. The algorithm allows for markers to be “pre-selected” by placement in the Selected Markers set upon initialization, and runs until the Selected Markers bin equals a certain maximum value. Probesets for the resulting selected markers are included in the fabrication of the custom microarray chip.

**Supplementary Figure 2a.**
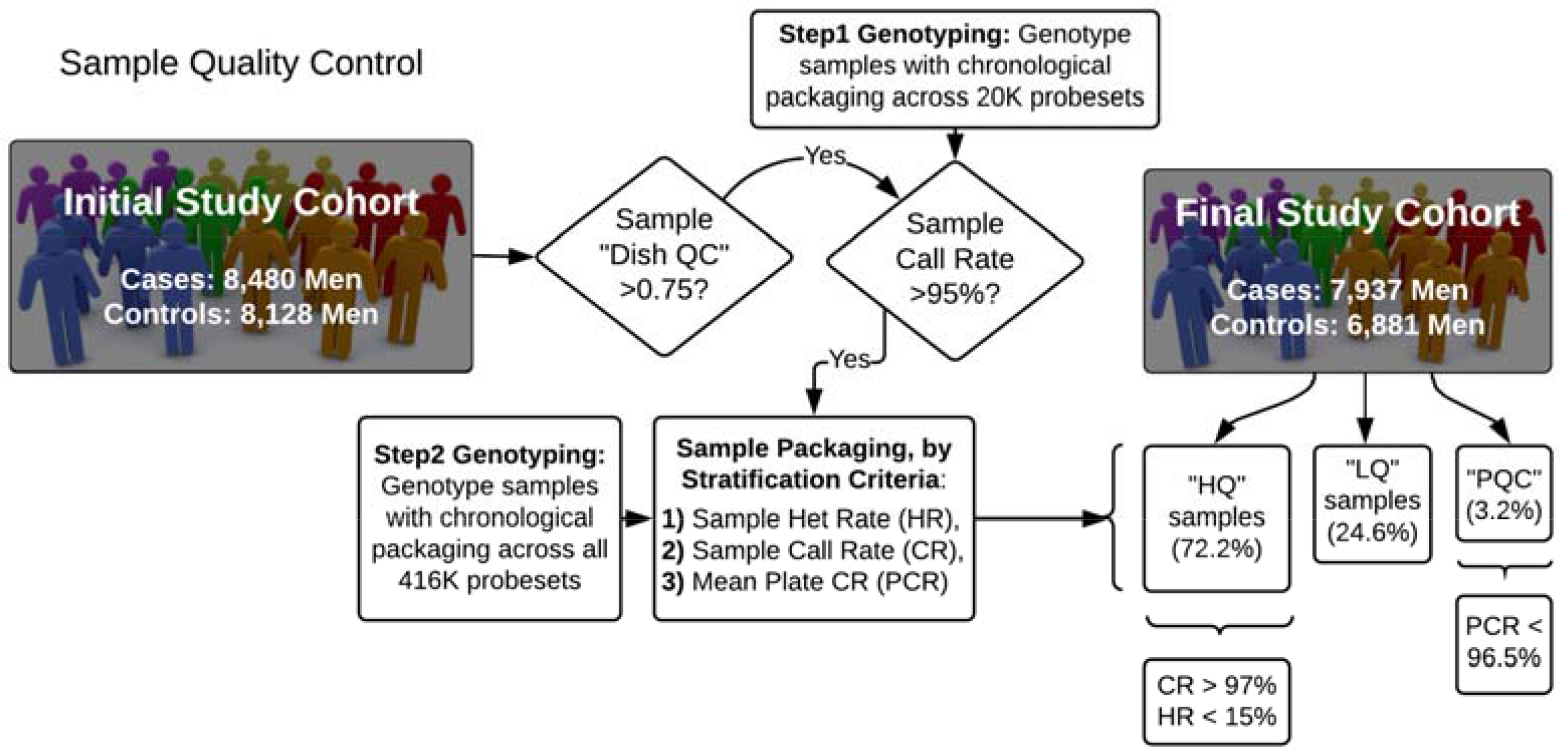
Sample Quality Control. Sample and Variant Quality Control Workflows. 2a. Sample Quality Control. 2b. Variant Quality Control.

**Supplementary Figure 2b.**
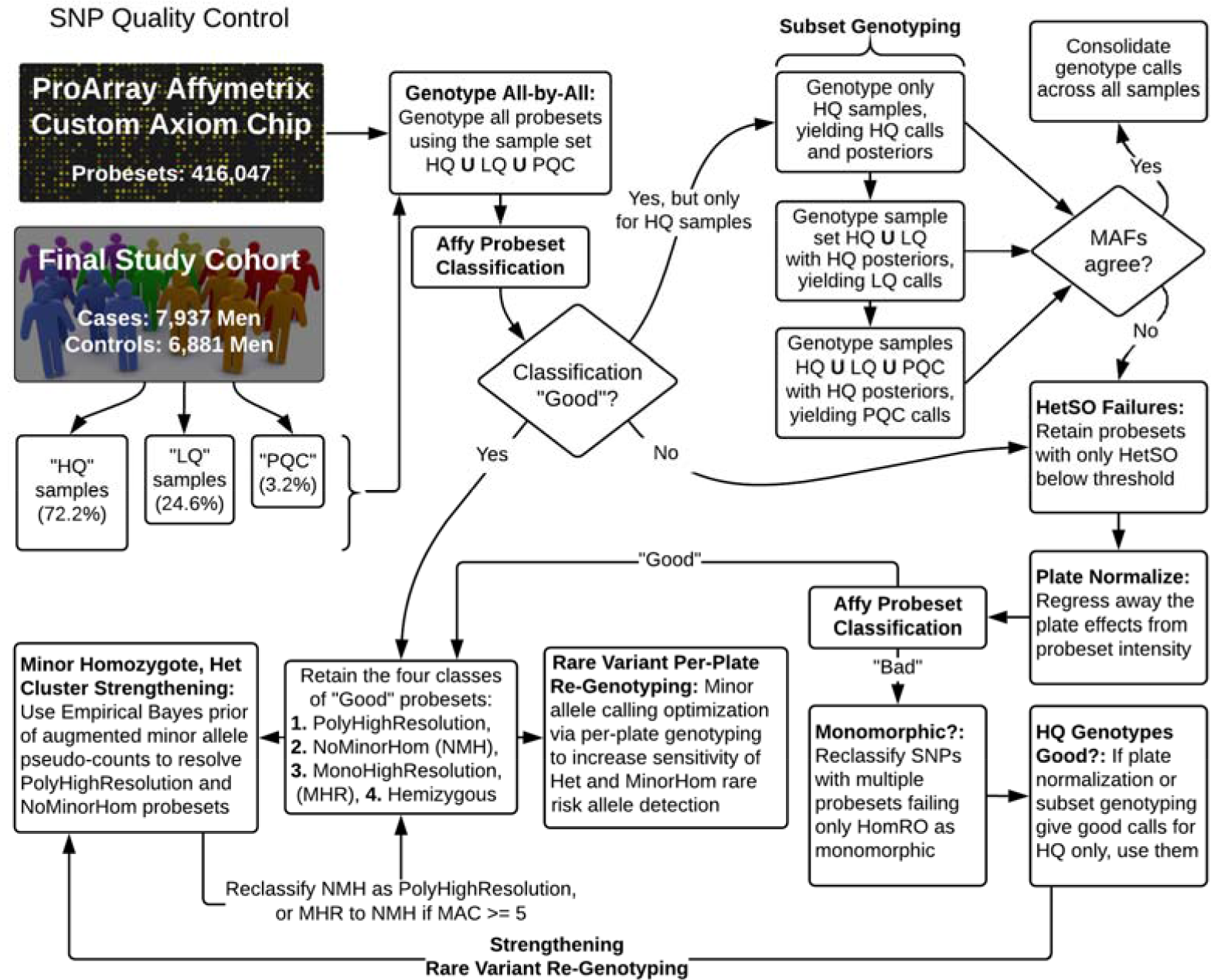
Variant Quality Control. Sample and Variant Quality Control Workflows. 2a. Sample Quality Control. 2b. Variant Quality Control.

**Supplementary Figure 3.**
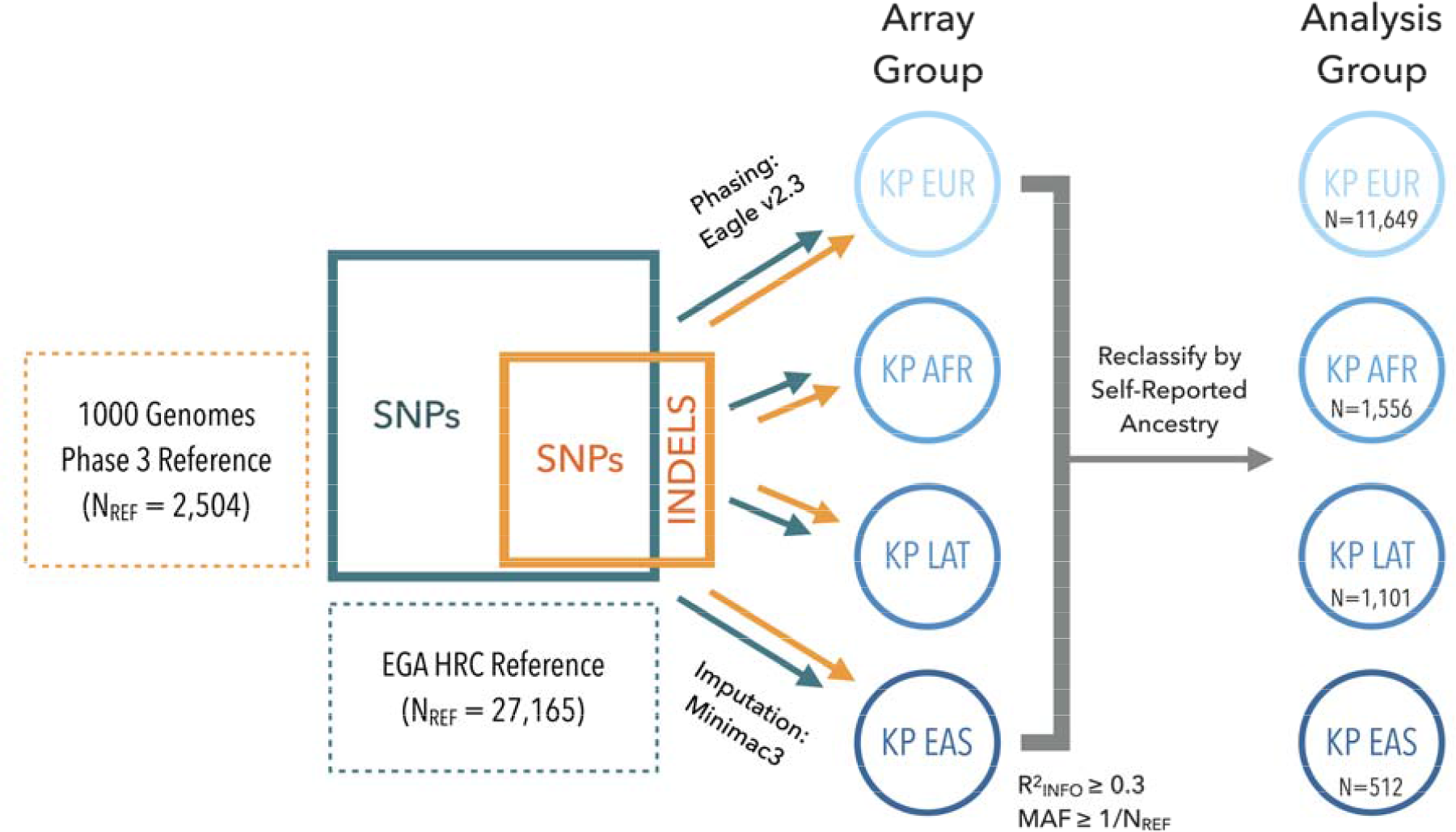
Genotype Imputation Workflow. Genotype Imputation Workflow. Depicted is the procedure implemented for imputing the Kaiser Permanente (KP) genotype data, from four ethnic groups: European ancestry (EUR), African ancestry (AFR), Latino ancestry (LAT), and East Asian ancestry (EAS). KP data were phased, reference-free (cohort-based), into haplotype-resolved genomes using Eagle v2.3. Next, single nucleotide polymorphisms (SNPs) were imputed using Minimac3 and a combined reference panel of Haplotype Reference Consortium (number of references, N_REF_: 27,165) and 1000 Genomes Project Phase III (number of references: 2,504) reference genomes. Furthermore, indel variants were imputed using the 1000 Genomes Project Phase III reference. Imputed SNPs and indels were combined, filtered based on imputation r^2^ (R^2^_INFO_) and minor allele frequency (MAF), and resegregated into analysis groups based on their self-reported ancestry (as opposed to the array groups with which they were genotyped).

**Supplementary Figure 4.**
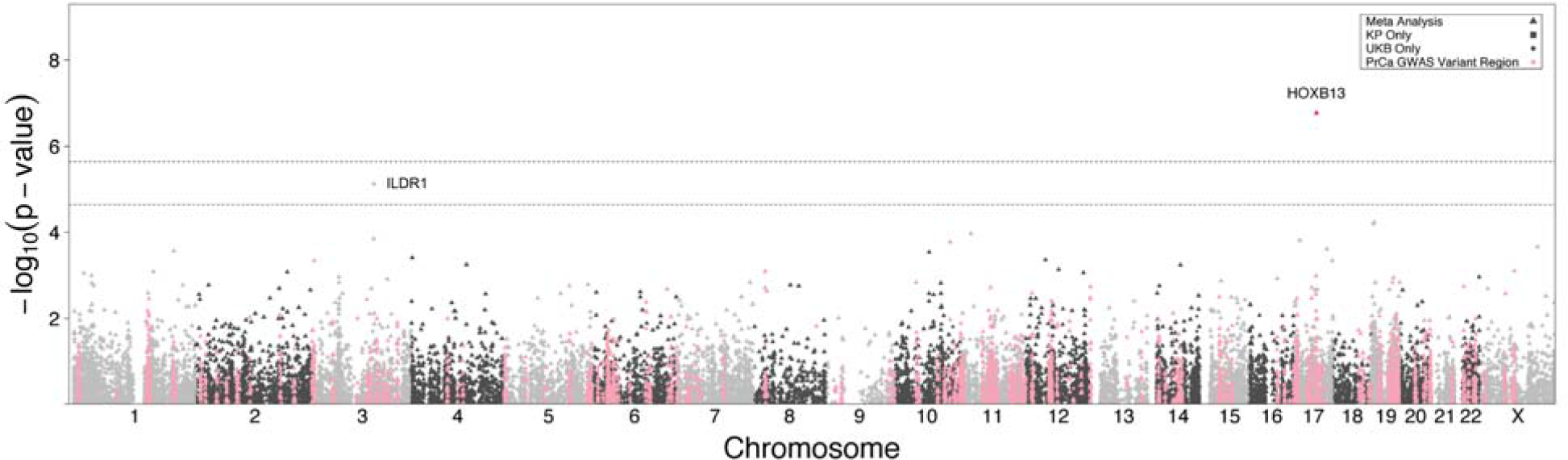
SKAT Gene-Based Rare Variant (MAF < 1%) Meta-Analysis of KP and UKB European-Ancestry Subjects. Gene-Based Test Manhattan Plot. Manhattan plot of associations for a gene-based meta-analysis between the Kaiser Permanente and UK Biobank. The associations (-log_10_(*P*-value), Y-axis) are plotted against the chromosome (1-22, X) and position (X-axis) of the modeled genes, with thresholds for Bonferroni-significant (*P* < 2.5*10^-6^) and suggestive (2.5*10^-5^ < *P* < 2.5*10^-6^) associations illustrated by dashed grey lines. Non-significant genes on odd and even chromosomes are colored in alternating shades. Triangular data points illustrate variants that were meta-analyzed between KP and UKB, while squares and circles indicate genes present exclusively in the KP or UKB summary statistics, respectively. Previously discovered PrCa loci are highlighted in pink for a 2 Mb window around the reported lead variant.

**Supplementary Figure 5a.**
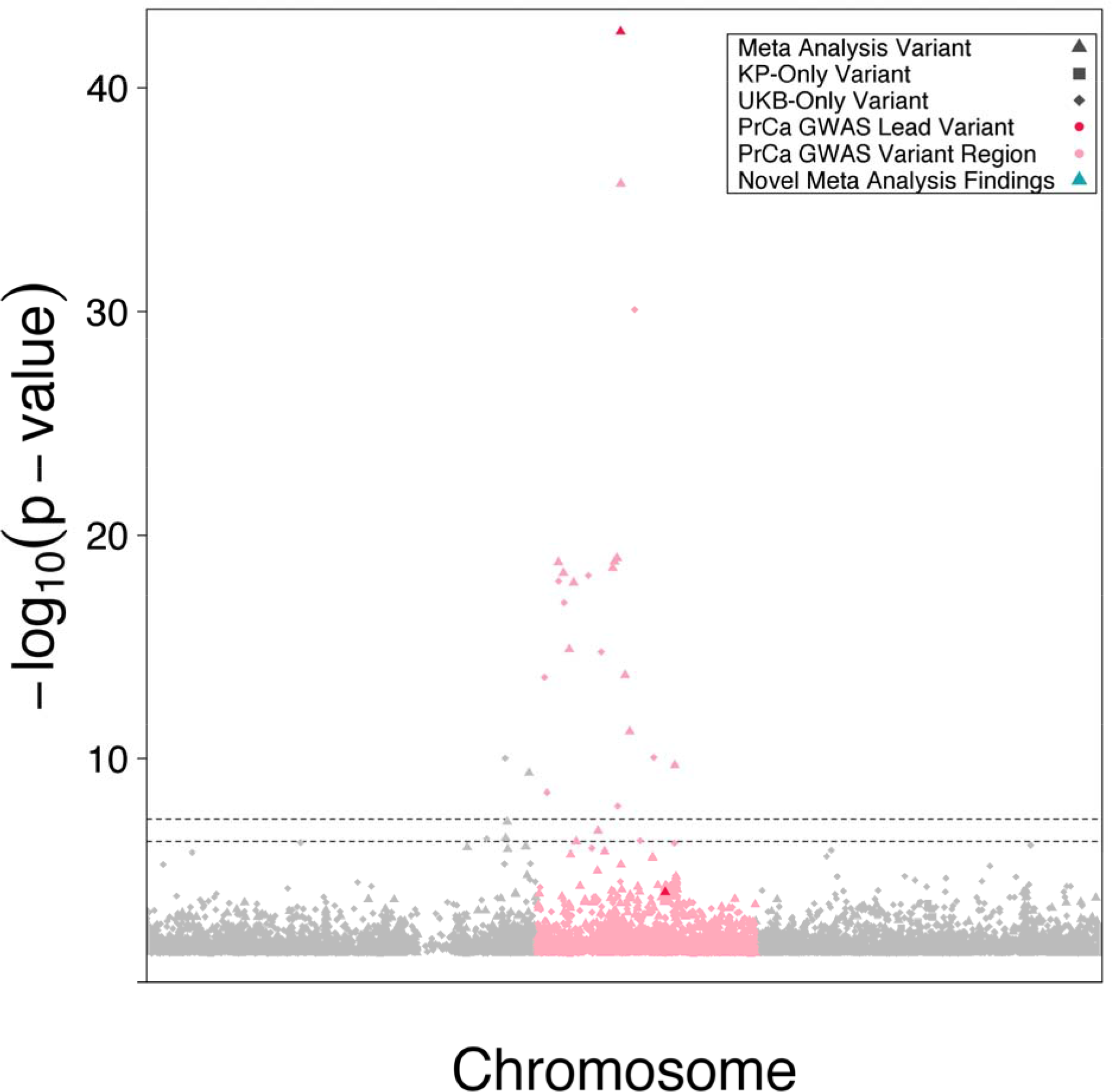
17q12 Locus Manhattan Plot. Associations and Linkage Disequilibrium at 17q12. S5a. Manhattan plot for meta-analysis of Kaiser Permanente and UK Biobank genotypes at the 17q12 locus, centered around the HOXB13 G84E missense variant rs138213197. Variants within 1Mb of the highly significant association at the rs138213197 SNP (*P* < 1*10^-40^) are colored in pink, demonstrating the width of the association peak. S5b. Linkage disequilibrium (LD) heatmap plot for all 17q12 variants with *P* < 5*10^-6^. Long range LD (beyond 1Mb) with respect to rs138213197 is illustrated.

**Supplementary Figure 5b.**
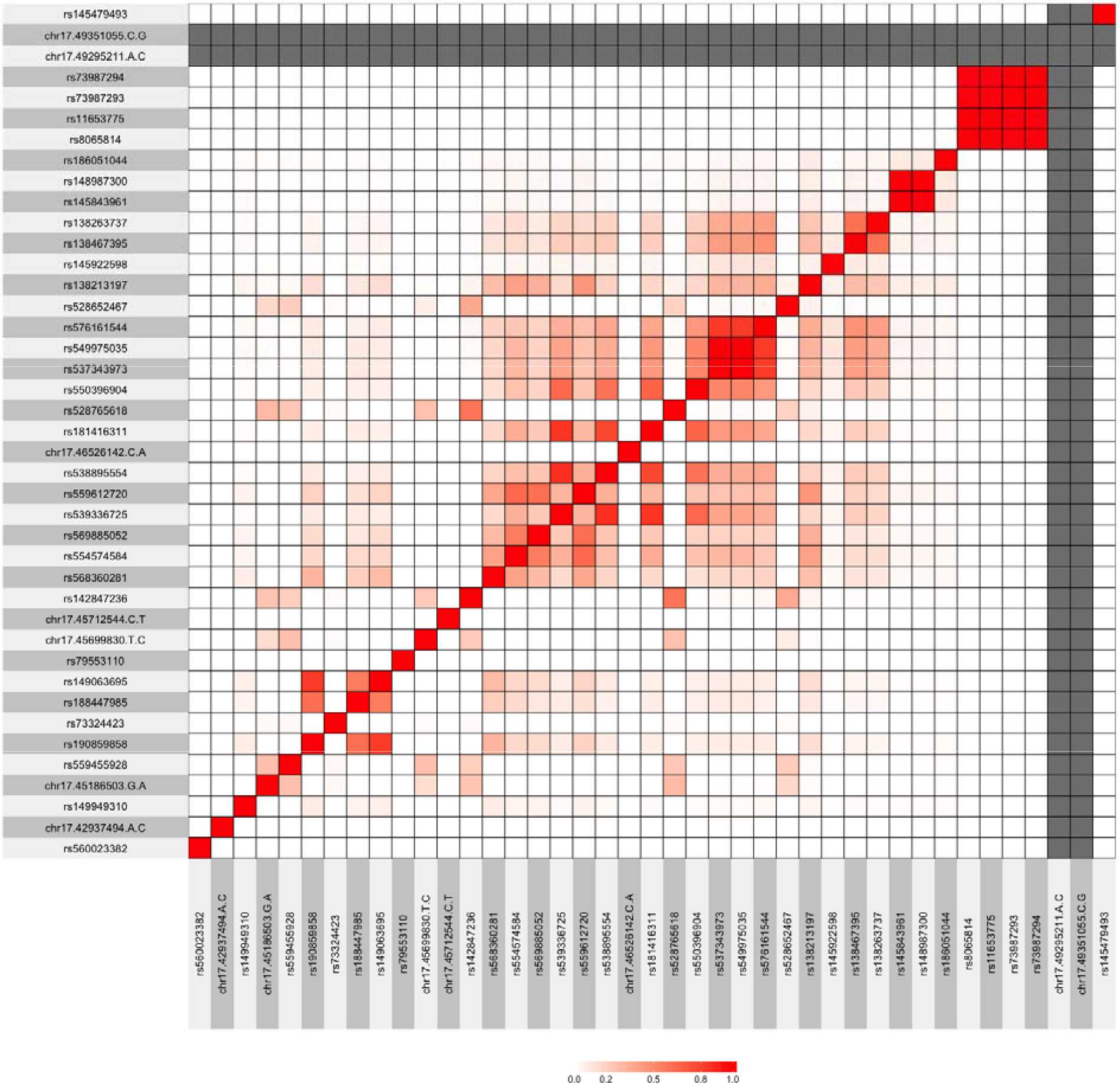
Linkage Disequilibrium at rare HOXB13 G84E missense variant rs138213197 (17q12) Associations and Linkage Disequilibrium at 17q12. S5a. Manhattan plot for meta-analysis of Kaiser Permanente and UK Biobank genotypes at the 17q12 locus, centered around the HOXB13 G84E missense variant rs138213197. Variants within 1Mb of the highly significant association at the rs138213197 SNP (*P* < 1*10^-40^) are colored in pink, demonstrating the width of the association peak. S5b. Linkage disequilibrium (LD) heatmap plot for all 17q12 variants with *P* < 5*10^-6^. Long range LD (beyond 1Mb) with respect to rs138213197 is illustrated.

**Supplementary Figure 6.**
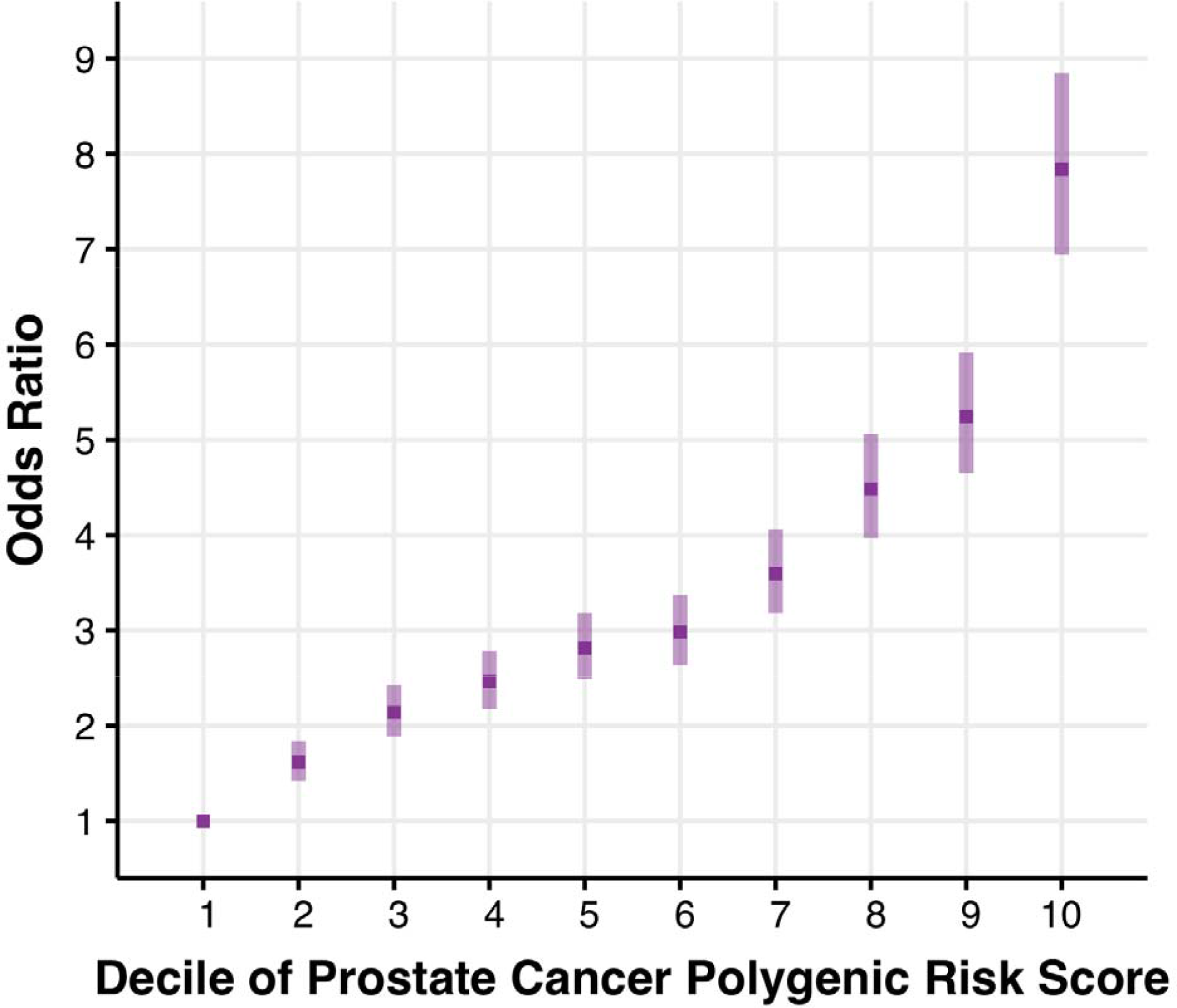
Polygenic Risk Score Modeling of Prostate Cancer Across KP and UKB Subjects “Prostate Cancer Polygenic Risk Score Performance. A polygenic risk score (PRS) of 187 previously reported prostate cancer (PrCa) risk variants was applied to subjects of European ancestry from two cohorts (Kaiser Permanente and UK Biobank). The Y-axis illustrates the magnitude of the odds ratio and 95% confidence interval for the association between PRS values and PrCa case-control status within a given decile of the PRS, in relation to the bottom decile as a reference group. Models were adjusted for age, body mass index, and principal components of ancestry.”

